# Revealing a coherent cell state landscape across single cell datasets with CONCORD

**DOI:** 10.1101/2025.03.13.643146

**Authors:** Qin Zhu, Zuzhi Jiang, Binyamin Zuckerman, Leor Weinberger, Matt Thomson, Zev Gartner

**Author notes:** **Author information:** Qin Zhu,; Zuzhi Jiang,; Binyamin Zuckerman,; Leor Weinberger,; Matt Thomson,; Zev J. Gartner.

## Abstract

Revealing the underlying cell-state landscape from single-cell data requires overcoming the critical obstacles of batch integration, denoising, and dimensionality reduction. Here, we present CONCORD, a unified framework that simultaneously addresses these challenges within a single self-supervised model. At its core, CONCORD implements a unified probabilistic sampling strategy that corrects batch effects via dataset-aware sampling and enhances biological resolution through hard-negative sampling. Remarkably, using only a minimalist neural network with a single hidden layer and contrastive learning, CONCORD surpasses state-of-the-art performance without relying on deep architectures, auxiliary losses, or external supervision. It seamlessly integrates data across batches, technologies, and even species to generate high-resolution cell atlases. The resulting latent representations are denoised and biologically meaningful—capturing gene co-expression programs, revealing detailed lineage trajectories, and preserving both local geometric relationships and global topological structures. We demonstrate CONCORD’s broad applicability across diverse datasets, establishing it as a general-purpose framework for learning unified, high-fidelity representations of cellular identity and dynamics.

## Introduction

Cells express thousands of genes to perform specialized functions and maintain homeostasis. Gene expression is highly correlated, orchestrated by intricate gene regulatory networks and cell-cell interactions that constrain cells to a structured, low-dimensional “state landscape” within the high-dimensional gene expression space^1,2^. Advances in single-cell technologies, particularly single-cell RNA sequencing (scRNA-seq), enable empirical mapping of this landscape. Emerging evidence suggests that such landscapes may contain diverse features—including discrete clusters, continuous trajectories, branching trees, and cyclic transitions—reflecting the underlying organization of cellular states^3,4^. However, the presence and arrangement of these features are typically unknown a priori, underscoring the need for computational methods that can robustly capture their topology and geometry to illuminate the principles of development, homeostasis, and disease progression.

Dimensionality reduction, a form of representation learning, is commonly employed to uncover the structure of the cell state landscape. By projecting high-dimensional data into a lower-dimensional space, key structural patterns become more tractable to visualize and analyze. However, conventional methods—such as principal component analysis (PCA), non-negative matrix factorization (NMF)^5^, and factor analysis^6^—often overemphasize broad cell type distinctions at the expense of subtle states, and can confound processes like differentiation with cell cycle progression. These challenges are exacerbated by batch effects— poorly understood sources of technical variation that obscure or skew genuine biological signals. Although an array of batch-correction tools—such as Harmony^7^, Scanorama^8^, Seurat^9^, scVI^10^, LIGER^11^ and MNN^12^ —have been developed, they frequently make strong assumptions about the structure of technical variation, leading to distortions from over- or under-correcting batch effects^13^. Furthermore, many face scalability issues when applied to massive atlas-level datasets.

Among emerging representation learning approaches, contrastive learning has recently shown promise for single-cell analysis^14–20^. Initially developed for domains such as image and natural language processing^21–23^, these methods learn informative cell representations by comparing similar (“positive”) cells against dissimilar (“negative”) ones within mini-batches—small subsets of cells iteratively sampled during training. However, current contrastive methods face fundamental limitations: supervised approaches require extensive manual annotation and struggle to generalize to novel states or continuous trajectories^19,20^, whereas unsupervised methods form mini-batches through uniform sampling, emphasizing coarse cell-type differences while overlooking subtle biological variation^14–17^. When applied across datasets, contrasting cells randomly sampled from different datasets can amplify dataset-specific artifacts rather than isolating true biological signals. While strategies involving generative adversarial networks (GANs)^17,24,25^, unsupervised domain adaptation via backpropagation^26^, and conditional variational autoencoders (CVAEs)^27^ attempt to mitigate batch effects, their objective of minimizing dataset-specific differences inherently conflicts with contrastive learning’s goal of maximizing differences between dissimilar cells, frequently leading to incomplete batch-effect correction and potentially introducing distortions to the latent space. This dilemma raises a critical question: can contrastive learning fully capture cellular diversity while minimizing batch effects?

Here, we address this open question by transforming a core limitation of contrastive learning—its sensitivity to mini-batch composition—into a strength. Our central insight is that mini-batch composition fundamentally determines the outcome of contrastive learning. We introduce CONCORD (COntrastive learNing for Cross-dOmain Reconciliation and Discovery), a framework that redefines the contrastive learning process through a probabilistic mini-batch sampling strategy combining dataset-aware sampling and hard-negative sampling. By strategically composing each mini-batch primarily with cells from the same dataset—thereby preventing the model from learning technical differences among batches while focusing on biological differences among cells—CONCORD simultaneously enhances embedding resolution and mitigates batch-specific artifacts. In contrast to prior methods that rely on complex architectures or auxiliary losses for batch correction, CONCORD achieves dimensionality reduction, denoising, and data integration solely through principled sampling. We demonstrate its effectiveness using a minimalist, single-hidden-layer neural network across simulated and real datasets spanning a range of biological and technical complexity. CONCORD consistently outperforms state-of-the-art methods, producing high-resolution, denoised encodings that robustly capture diverse structures—including clusters, loops, trajectories and trees—reflecting bona fide biological processes even when the data originate from multiple technologies, time points, or species. This versatile framework scales from small to large datasets, generalizes to modalities beyond scRNA-seq, and establishes a rigorous foundation for next-generation single-cell machine learning models to drive diverse downstream biological discoveries.

## Results

### The CONCORD framework

Analyses of single-cell sequencing data suggest that gene expression is not randomly sampled; rather, the mechanism of gene regulation imposes strong constraints, producing dynamically changing gene co-expression patterns reflected as intricate structures in the low dimensional embedding of cells^1–3,28^. For example, at homeostasis, cells typically form discrete clusters corresponding to stable types or states, with adjacent clusters representing closely related states (Figure 1A, left). In developmental or pathological contexts—such as early embryogenesis, tissue repair, or tumorigenesis—cells often follow branching trajectories from progenitors to terminal fates, with semi-stable intermediate states forming denser clusters (Figure 1A, middle). Cyclic gene expression programs, such as those regulating the cell cycle, give rise to loop-like structures^3,4^ (Figure 1A, right). Despite these rich patterns, conventional dimensionality reduction methods like PCA or NMF capture only partial representations of the cell state landscape, either oversimplifying complex structures or disproportionately emphasizing certain features while obscuring others.

**Figure 1.**
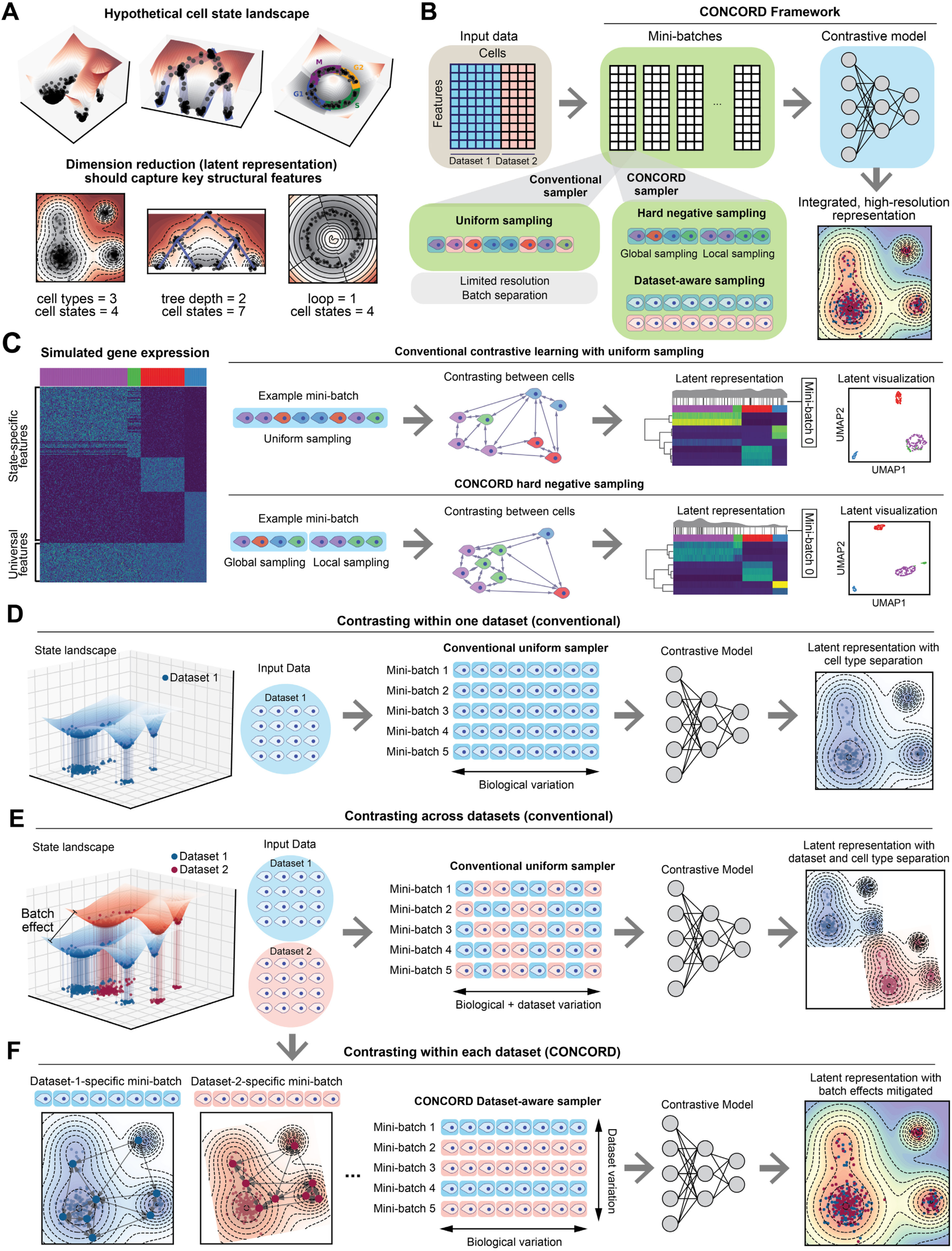
CONCORD mini-batch sampling enables high-resolution, batch-effect-mitigated representation learning of single cell data. (**A**) Schematic of hypothetical cell-state landscapes and corresponding low-dimensional representations that capture key structural features. (**B**) Overview of the CONCORD framework, which replaces the conventional mini-batch sampler with a joint hard-negative and dataset-aware sampling scheme, enabling integrated, high-resolution representation learning with a minimalist contrastive model. (**C**) Uniform versus hard-negative sampling in a simulated four-state dataset. Heatmaps show simulated expression and latent space, accompanied by density curves with black lines indicating the distribution of cells in an example mini-batch under each scheme. Resulting UMAP embeddings are shown. (**D**) Contrastive learning on a single dataset using the conventional uniform sampler, which draws cells uniformly from the entire dataset to form mini-batches. (**E**) Applying standard contrastive learning across multiple datasets mixes datasets within mini-batches and amplifies batch effects, as reflected in the latent embeddings. (**F**) CONCORD mitigates batch effects by predominantly contrasting cells within each dataset and randomly shuffling mini-batches across epochs.

We hypothesized that a representation-learning approach capable of encoding cells based on gene co-expression programs would provide a more comprehensive view of the cell state landscape. Recent evidence suggests that self-supervised contrastive learning recovers sparse, structured gene co-expression signals from high-dimensional data^29^, significantly improving clustering and cell-type classification performance^15–17^ (see Methods). Like many modern machine learning methods, contrastive learning relies on mini-batches— small subsets of data sampled iteratively during stochastic gradient descent—as the basic unit of training. However, contrastive learning is uniquely sensitive to mini-batch composition: each cell is contrasted against every other cell within the same mini-batch, making the mini-batch the universe over which learning is defined. By differentiating each cell from others in the mini-batch, the model learns features that distinguish distinct cellular states. Simultaneously, aligning augmented versions of the same cell (typically generated through random masking) encourages the model to capture robust gene co-expression patterns, rather than relying on the expression of individual genes^29^. As a result, the learned representations are intrinsically more robust to technical noise and dropout—pervasive artifacts in single-cell datasets^30^.

This reliance on within-mini-batch comparisons makes the sampling strategy— which dictates mini-batch composition—a critical determinant of the learned representation^31^. Existing methods typically adopt uniform sampling across single-cell datasets (Fig. 1B), leading to two key limitations. First, uniform sampling emphasizes broad differences (e.g., major cell types) while underrepresenting rare subpopulations or subtle distinctions, resulting in poor resolution of fine-scale cellular states (Figure 1C). Second, mixing cells from different datasets within the same mini-batch can amplify technical dataset-specific differences, also known as “batch effects,” inadvertently encoding these batch effects rather than isolating biologically meaningful variation.

To address the first issue, we adopt hard-negative sampling^32^, in which each mini-batch is enriched with closely related cells (Fig. 1C), encouraging the model to extract features that distinguish these “hard negatives”. We implemented two variants. First, inspired by previous work^31^, we developed a k-nearest neighbor (kNN)-based sampler that probabilistically draws cells from both their local neighborhoods and the global distribution. Local sampling—guided by a coarse graph approximation of the cellular state landscape—compels the model to contrast each cell with its neighbors, enabling detection of subtle differences between closely related states. Simultaneously, global sampling preserves a broad perspective of major cell types, ensuring robust encoding of large-scale distinctions. By iteratively presenting the model with local neighborhoods (e.g., T cells in one mini-batch, epithelial cells in another) alongside the global distribution, the model allocates capacity to represent both large-scale distinctions and nuanced local details, leading to improved resolution in the learned latent space (Figure 1C). As a second variant (referred to as *hcl* mode, following the original authors), we implement the approach by Robinson et al.^32^, which uses Monte-Carlo importance sampling to approximate the expected loss of hard-negative sampling without explicit neighborhood-based sampling (see Methods).

When applied to a single dataset, contrastive learning effectively captures biological variation in the latent space (Figure 1D). However, with uniform sampling across multiple datasets, both biological and dataset-specific variations are encoded, yielding latent spaces that separate by dataset as well as cell type (Figure 1E). To address this, we introduce a dataset-aware sampler that restricts mini-batches to a single dataset, ensuring contrasts reflect only biological differences—as in the single-dataset setting (Figure 1F). Dataset-specific biases are further diminished through random mini-batch shuffling: if such signals are encoded in one batch, they are disrupted and overwritten by subsequent mini-batches from other datasets. Consequently, only biologically meaningful signals, such as gene co-expression patterns, persist throughout training, producing latent spaces that reflect biological variation with minimal batch effects (Figure 1F). In cases where datasets have minimal or no overlap, a leaky dataset-aware sampler enables soft alignment without imposing artificial harmonization, supporting flexible integration that respects dataset-specific signals (Supplemental Figure 1A). Notably, this approach does not perform any explicit modeling of batch effects; instead, it selectively captures and encodes biological programs shared across datasets. Unlike prior batch-correction strategies that struggle in contrastive settings due to competing objectives, CONCORD integrates batch correction directly into the contrastive learning process via its sampling design, producing latent representations inherently robust to batch effects.

Both the hard-negative and dataset-aware samplers follow a unified principle: probabilistically structuring mini-batches to balance global biological diversity with local and dataset-specific variation. We integrate both samplers into a joint sampling framework, where the likelihood of selecting a cell satisfies both sampling schemes (Supplemental Figure 1A, B, also see Methods). This generalized sampling strategy fundamentally reconfigures contrastive learning, enabling high-resolution representation learning and robust dataset integration within a single contrastive objective, and forms the core of the CONCORD framework (Supplemental Figure 1C). With this simple innovation, CONCORD outperforms state-of-the-art methods using only a minimalist encoder with a single hidden layer, demonstrating that sampling design alone can transform contrastive learning performance on single cell data—even without deep or complex architectures. This simplicity reduces training data requirements, enhances robustness, and increases interpretability of the learned latent space.

### CONCORD learns denoised latent representations that preserve underlying structures

Recovering biologically meaningful insights from single-cell data requires preserving the underlying geometric and topological structure of the gene expression space. To evaluate whether CONCORD meets this criterion, we benchmarked its performance on a suite of simulated datasets. As existing simulators often fail to generate complex biological structures like branching or loops, we developed a custom workflow to create realistic structures with flexible control over noise and batch effects (Figure 2A).

**Figure 2.**
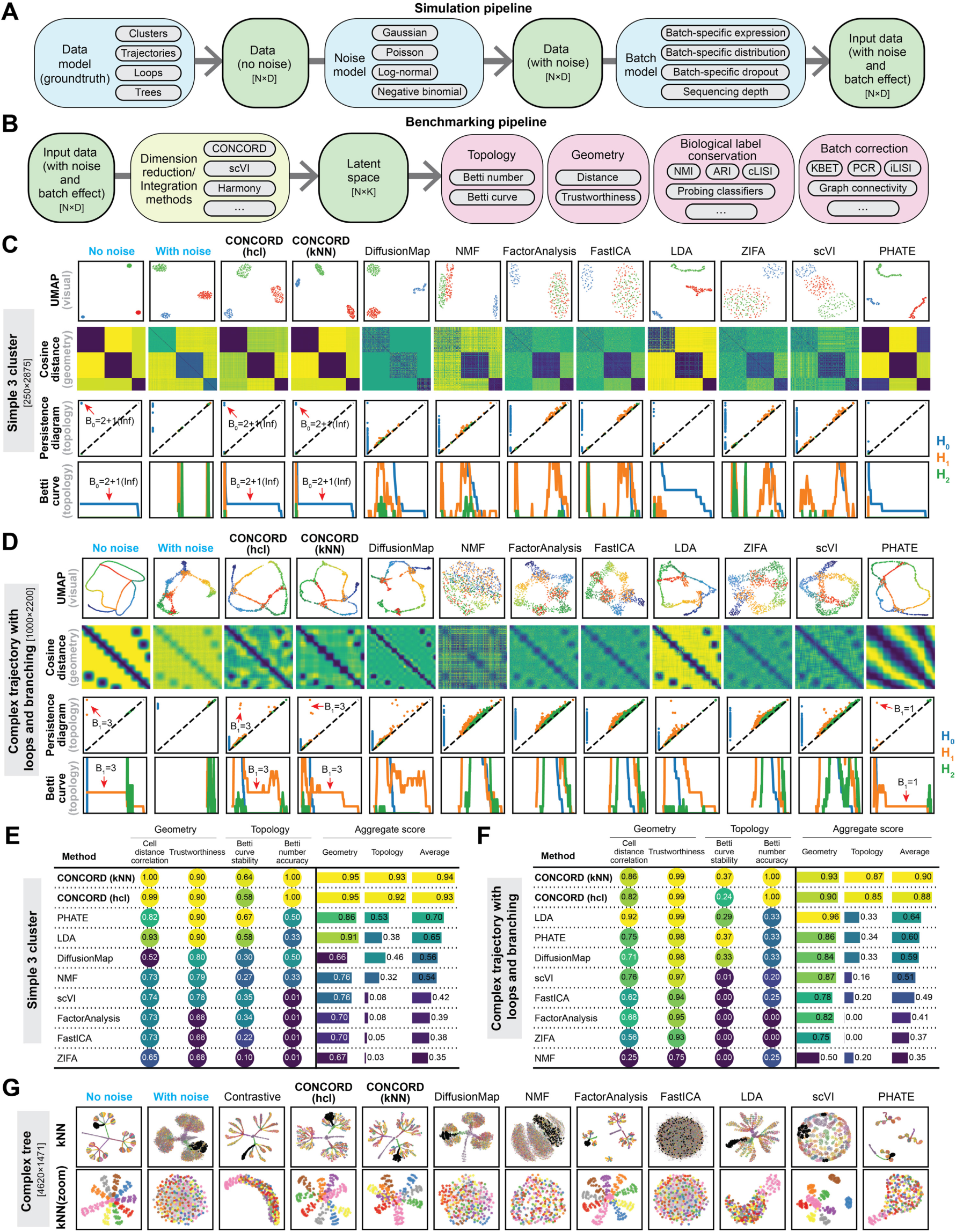
Benchmarking CONCORD and other dimensionality-reduction methods across diverse structures. **(A)** Schematic of the simulation pipeline, which first produces a noise-free gene expression matrix based on a user-defined data structure, then introduces noise following a specified noise model, and finally applies batch effects. (**B**) Schematic of the benchmarking pipeline. Latent representations from each method are compared with the noise-free ground truth to assess preservation of topological and geometric features. The scIB^33^ package and probing classifiers are used to evaluate biological label conservation and batch harmonization. (**C**) Performance of CONCORD and competing methods on a three-cluster simulation with dimensions listed. UMAP embeddings, cosine-distance matrices and persistent-homology analysis (persistence diagram and Betti curves) are shown for each method. The H_0_ point at infinity was excluded from the persistence diagram and curve. (**D**) Performance on a complex trajectory with three loops, highlighting the same diagnostic plots as in (C). (**E, F**) Summary of key geometric and topological performance metrics for the cluster simulation (**E**) and the complex trajectory simulation (**F**). (**G**) KNN-graph visualization of the latent spaces from a complex-tree simulation, with zoomed-in views of the highlighted branch.

To assess the quality of learned representations, we established a comprehensive evaluation pipeline. While standard benchmarks like the scIB framework^33^ effectively measure label preservation and batch mixing, they are often insufficient for evaluating the preservation of complex biological structures^34,35^. We therefore supplemented them with probing classifiers^36,37^—a standard approach for evaluating representation learning—to assess the conservation of biological labels in the latent space. Additionally, to quantify structure fidelity, we incorporated geometric metrics like trustworthiness and global distance correlation, as well as topological data analysis (TDA) based on persistent homology and Betti numbers (Figure 2B). These metrics evaluate embedding at complementary scales: trustworthiness quantifies local neighborhood preservation, while persistent homology captures global topological features—such as clusters (Betti-0), loops (Betti-1), and voids (Betti-2). These features are visualized in persistence diagrams and Betti curves, where stable structures appear as long-lived features in the persistence diagram and extended plateaus in the Betti curve, whereas transient, noise-induced features vanish quickly.

We evaluated both CONCORD variants on a simple, single-batch simulation consisting of three well-separated clusters corrupted by cluster-specific Gaussian noise (Figure 2C, Supplemental Figure 2A). Compared to a broad set of dimensionality reduction methods—including diffusion map^38^, NMF^39^, Factor Analysis^40^, FastICA^41^, Latent Dirichlet Allocation (LDA)^42^, ZIFA^43^, scVI^10^, and PHATE^44^—CONCORD cleanly separated clusters, as reflected in both the latent space and pairwise distance matrices. In contrast, many methods failed to fully resolve the clusters or introduced spurious structures, such as trajectory-like artifacts (Figure 2C). Persistent homology confirmed these observations: CONCORD’s Betti-0 plateau accurately reflected the expected three-cluster topology and closely matched the noise-free reference, highlighting its strength in both denoising and structure preservation.

On a more complex simulation with three loops and multiple branching points (Figure 2D, Supplemental Fig. 2B), CONCORD was the only method that accurately recovered the full topology. Other methods either distorted the structure or failed to detect the correct number of loops in Betti analysis, likely due to excessive noise retention. Although PHATE produced a visually similar embedding, its Betti curve identified only a single persistent loop, indicating that critical topological features were obscured in its latent space.

Quantitative evaluation of geometric and topological metrics confirmed that CONCORD consistently outperformed competing methods (Figure 2E, 2F). Notably, CONCORD maintained high trustworthiness across a wide range of neighborhood sizes, underscoring its ability to preserve local geometry at multiple scales (Supplemental Figure 2C, 2D). In contrast, other methods exhibit considerable declines in trustworthiness, indicating a loss of fine-scale geometric relationships.

To assess the impact of hard-negative sampling, we simulated a hierarchical branching tree (Figure 2G, Supplemental Figure 2E-G). Without hard negative sampling, sub-branches were unresolved. Moderate enrichment of hard negatives substantially improved resolution for both CONCORD variants, with the kNN mode being more susceptible to excessive local focus, which obscured global distinctions (Supplemental Figure 2F, G).

### CONCORD learns a coherent, batch-effect-mitigated latent representation

Batch effects often appear as dataset-specific global signals that can obscure biological variation. In CONCORD, these signals rapidly diminish during training when mini-batches are restricted to single datasets (Figure 1E, Figure 3A). Unlike conventional batch-correction methods that rely on explicit alignment models, CONCORD makes minimal assumptions about the source or form of batch effects and instead prioritizes learning coherent, biologically meaningful gene covariation patterns. This leads to more accurate preservation of biological structure while mitigating technical artifacts.

**Figure 3.**
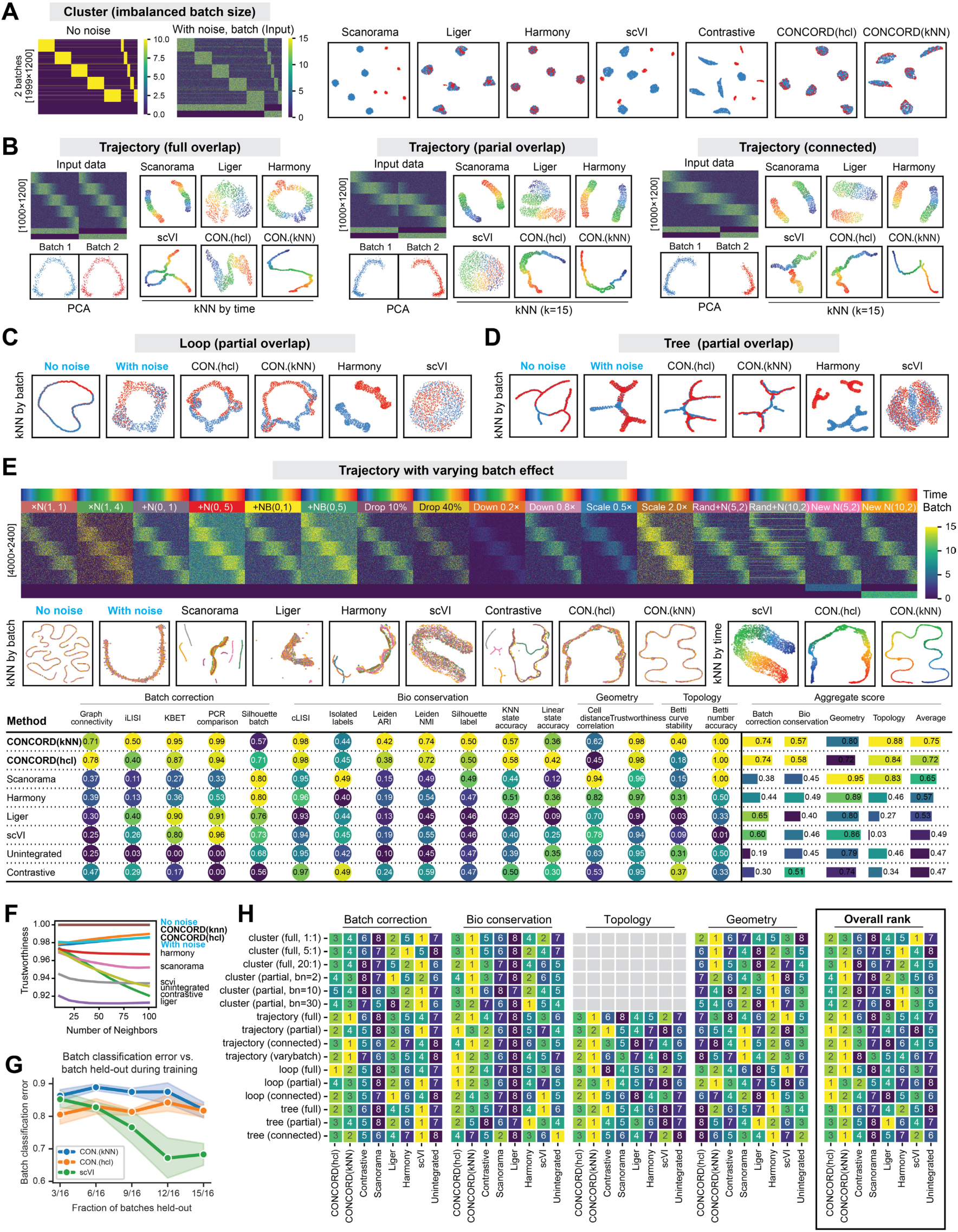
Benchmarking CONCORD and other data-integration methods across diverse structures. (**A**) Two-batch, five-cluster simulation with imbalanced batch sizes. Heatmaps show the noise-free ground truth and the input data with noise and batch effects. Latent spaces from each method are visualized by UMAPs, colored by batch. Full cluster-simulation results are in Supplemental Figure 3A-B. “Contrastive” refers to naïve contrastive learning that uses the same encoder architecture and objective as CONCORD, but with uniform sampling. (**B**) Trajectory simulation with varying batch overlap. The input structure is shown by a heatmap and PCA. For each method, the latent space is visualized by a kNN graph (k = 15) colored by simulated time to assess cross-batch integration along the trajectory. (**C**) Loop simulation with varying batch overlap. kNN graphs are shown for the ground truth (edges omitted) and for CONCORD and selected methods. Full results are in Supplemental Figure 3C. (**D**) Tree simulation with varying batch overlap. kNN graphs are shown for the ground truth, CONCORD, and selected methods. Full results are in Supplemental Figure 3D. (**E**) Trajectory simulation with 16 batches, each with a different batch effect (heatmap). kNN graphs (k = 15) colored by batch are shown for each method’s latent embeddings; for scVI and both CONCORD modes (*hcl* and *kNN*), kNN graphs colored by simulated time are also shown. A table displaying detailed benchmarking metrics is provided; see Methods for details on metric definitions. (**F**) Trustworthiness across neighborhood sizes for the multi-batch simulation in (E). (**G**) Prediction with limited training data for scVI and CONCORD. A specified number of batches were held out during training; we ran five replicates with random batch withholding and quantified batch mixing via kNN-based batch classification error (k = 30). Means and 95% confidence intervals are plotted. (**H**) Ranking of integration methods across simulated data, showing ranks for batch correction, biological-label conservation, topological and geometric metrics, and overall score. For cluster simulations, Betti curves become noisy when the number of clusters exceeds three, and we did not find a robust way to infer Betti numbers; therefore, topology scores were excluded for these datasets.

We first evaluated CONCORD on a simulated five-cluster dataset with varying noise, batch effects, and batch size imbalance (Figure 3A, Supplemental Figure 3A). Across these conditions, CONCORD was the only method to robustly recover all five clusters. This success is attributable to its dataset-aware sampler, as using a conventional uniform sampler (i.e., the naïve contrastive approach) resulted in pronounced batch effects. In more challenging scenarios with more batches and greater imbalance, CONCORD and Harmony were the only methods that consistently separated the underlying clusters (Supplemental Fig. 3B).

Single-cell studies often involve continuous state transitions sampled across different conditions, where cell states may only partially overlap. Methods that make explicit assumptions about the data structure—such as requiring matched clusters—often fail in these scenarios and produce distorted embeddings. We systematically tested this by simulating batch effects on trajectories, loops, and trees with varying degrees of state overlap (Figure 3B-D, Supplemental Figure 3B-D). Many competing methods exhibited poor alignment and introduced artificial structures. In contrast, both CONCORD variants consistently recovered the correct topology with reduced noise, even when the overlap between batches was minimal.

We further tested performance on a trajectory with 16 distinct batch effects (Figure 3E). While scVI and CONCORD both aligned the batches, scVI showed incomplete alignment at fine resolution. In contrast, CONCORD—particularly the *kNN* variant—achieved superior alignment and noise reduction. Quantitative metrics confirmed these observations: CONCORD preserved local geometry, evidenced by high trustworthiness (Figure 3E,F), while exhibiting lower global distance correlation—a common trade-off in manifold learning^45,46^. Robustness was further demonstrated in a stress test where models were trained on a few randomly selected batches and used to predict the remaining ones (Figure 3G). CONCORD maintained strong alignment, whereas scVI’s performance degraded markedly as the number of training batches decreased. This suggests CONCORD’s robustness stems from learning gene co-expression programs rather than explicitly modeling and correcting batch effects.

Across all simulations, CONCORD achieved high biological label conservation (Figure 3G, Supplemental Table 1), with slightly lower batch-correction scores because it does not explicitly merge batches. By contrast, although scVI achieved high batch-mixing scores, it often produced over-mixed embeddings that obscured underlying structure (Supplemental Figure 3). The aggregate geometric score for CONCORD was reduced by its lower global distance correlation despite consistently strong trustworthiness; however, for data with manifold structures—such as single cell data—global distances are often not reflective of true distance relationships between cell states. Therefore, preserving local neighborhood fidelity is typically prioritized in single cell analysis^47^.

Nevertheless, CONCORD consistently ranks among the top methods for topological preservation, biological label conservation, and overall performance. These results demonstrate that CONCORD provides a reliable and generalizable framework for dimensionality reduction and batch correction, even when the data structure is unknown or batch overlap is limited.

### CONCORD aligns whole-organism developmental atlases and resolves high-resolution lineage trajectories

To assess whether CONCORD captures biologically meaningful structures, we benchmarked it against popular integration methods on *C. elegans* embryogenesis^48^—a well-characterized system with a nearly invariant lineage tree^49^ that is also conserved in the related species *C. briggsae*^50^. Packer et al. initially generated a lineage-resolved atlas of *C. elegans*^48^, which was recently expanded by Large et al. to include over 200,000 *C. elegans* cells and 190,000 *C. briggsae* cells^50^. With expert-curated annotations generated through iterative, labor-intensive zoom-in analyses and validated by fluorescent imaging, these datasets provide an ideal benchmark for evaluating whether integration methods can accurately reconstruct and align developmental trajectories across species.

We first tested CONCORD on the original *C. elegans* atlas (>90,000 cells). The resulting embedding revealed disconnected trajectories among early-stage cells, which we hypothesized reflected missing states. These gaps persisted even after including *C. elegans* cells from the expanded Large et al. dataset. We therefore collected a new *C. elegans* dataset enriched for early embryos; adding this dataset resolved the gaps and yielded a continuous trajectory from zygote to terminal fates (Supplemental Fig. 4A). Using the extensive cell-type and lineage annotations, we benchmarked CONCORD against other methods for batch correction and label conservation, and assessed its sensitivity to key hyperparameters (Supplemental Figure 4B-E). CONCORD significantly outperformed existing methods, with stable performance across the recommended hyperparameter range. Notably, the effect of hard-negative sampling mirrored trends observed in simulations: moderate local enrichment improved resolution, whereas excessive local sampling disrupted global structure (Supplemental Figure 4F).

When applied to over 410,000 cells from the combined cross-species dataset and our new early-embryo collection, CONCORD generated a unified developmental atlas that closely matched the expert annotations, achieving cross-species alignment and resolving lineages at ultra-high resolution (Figure 4A,B). Both the *hcl* and *kNN* modes yielded similar, high-quality embeddings (Figure 4A, Supplemental Figure 5A). Because scIB^33^ could not scale to this dataset, we quantified integration performance using probing classifiers to assess batch mixing, cell type, and lineage label preservation (Figure 4C).

**Figure 4.**
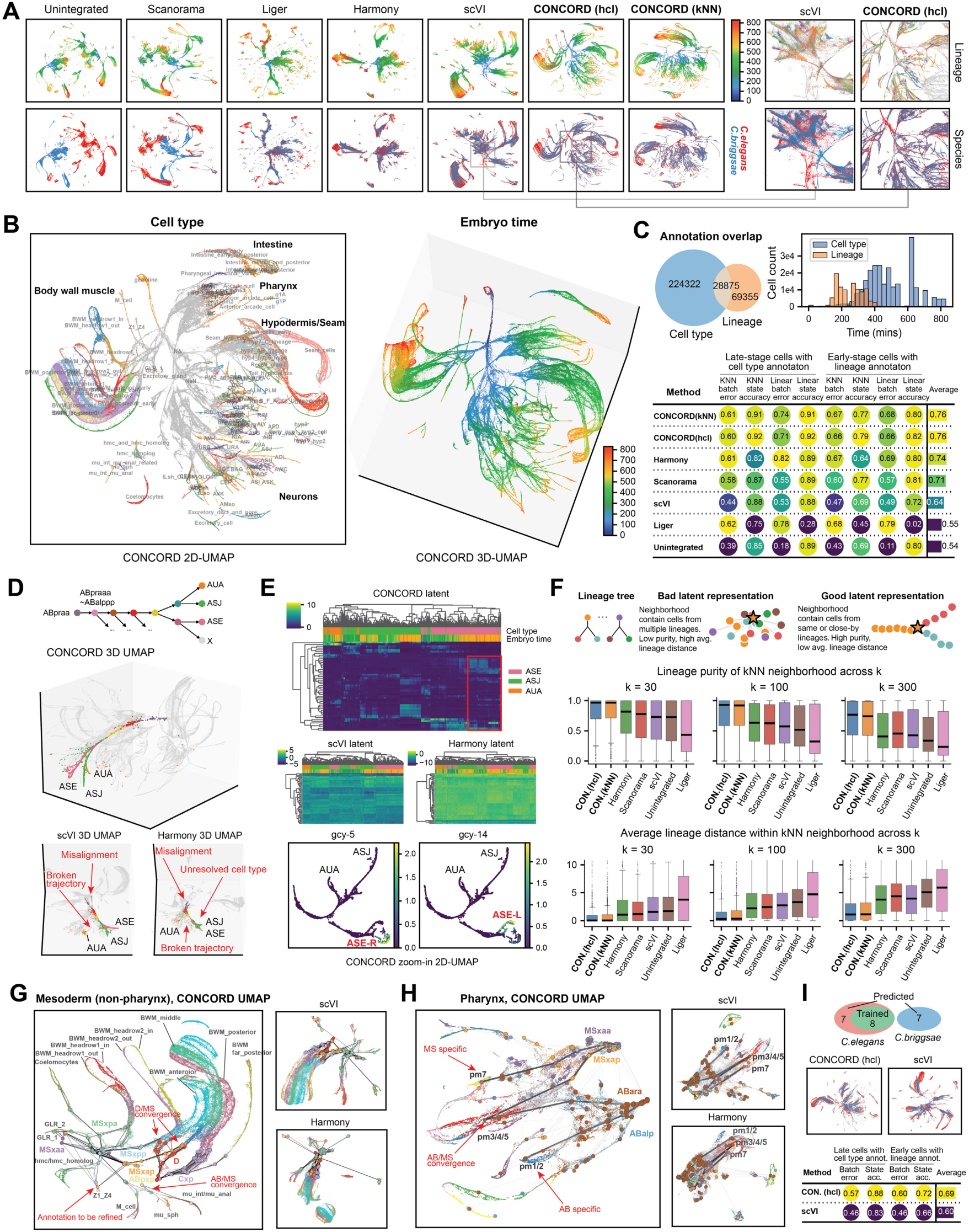
Benchmarking CONCORD on *C. elegans/C.briggsae* embryogenesis atlas. **(A)** UMAPs from CONCORD and other integration methods, colored by inferred embryo time and species. Zoomed-in UMAPs for scVI and CONCORD (*hcl*) show approximately matched regions, colored by lineage and species. (**B**) Global 2D and 3D CONCORD (*hcl*) embeddings colored by cell type and inferred embryo time. (**C**) Overlap between expert-curated cell-type and lineage annotations. A histogram shows lineage annotations concentrated in early-stage cells and cell-type annotations predominantly in late-stage cells. Integration performance was evaluated separately for early-stage cells (lineage labels) and late-stage cells (cell-type labels) using probing classifiers. (**D**) Global 3D UMAPs of CONCORD, scVI and Harmony, highlighting cells mapped to the lineage sub-tree that give rise to ASE, ASJ and AUA neurons. For each method, the most representative view was selected. (**E**) Heatmap showing the top 50 most variable latent dimensions in the ASE, ASJ, and AUA neuron subset for scVI, Harmony, and CONCORD (*hcl*). Expression of *gcy-5* and *gcy-14* is overlaid on a zoomed UMAP recomputed from the CONCORD latent space. (**F**) Lineage purity and average lineage distance computed across 2,000 randomly selected kNN neighborhoods for each method. For each randomly sampled anchor cell, we retrieve its k nearest neighbors in the embedding and compare their lineage relationships to the lineage graph. Purity is the fraction of neighbors assigned to the same lineage as the anchor; average lineage distance is the mean hop distance on the lineage tree from the anchor to its neighbors. Box plots show: median (center line), quartiles (box limits), 1.5× IQR (whiskers), and outliers (points). (**G**) Zoomed-in UMAPs for mesoderm (excluding pharynx), highlighting major input lineages and cell types. Each lineage is represented by its cluster medoid; edges connect parental lineages to daughters following the lineage tree. (**H**) Zoomed-in UMAPs for pharynx, annotated by cell type and broad input lineages. Selected lineage paths yielding pm1/2, pm3–5, and pm7 are highlighted. (**I**) scVI and CONCORD were trained on the combined *C. elegans* data from Packer et al.^48^ and our newly-collected batch, then used to project the full atlas including *C. elegans* and *C. briggsae* data from Large et al^50^. Resulting UMAPs are colored by species, and integration performance was evaluated with probing classifiers.

CONCORD excelled on these metrics, whereas other methods either failed to fully align the species or lost resolution, consistent with visual inspection of the UMAP embeddings. As the complexity of the learned structure exceeded the capacity of 2D UMAP, we encourage readers to explore the interactive 3D visualizations (https://qinzhu.github.io/Concord_documentation/galleries/cbce_show/#tabbed_1_1).

Projecting the lineage tree onto CONCORD’s embedding revealed strong concordance with established lineage and fate relationships (Supplemental Figure 5B). For example, the ASE, ASJ, and AUA neurons—derived from AB progenitors—formed branching trajectories that mirrored their true lineage structure (Figure 4D). In contrast, other methods introduced discontinuities, failed to resolve key bifurcations, or generated artificial structures. Strikingly, CONCORD’s latent space resolved ASE-left (ASEL) and ASE-right (ASER) neurons, characterized by differential expression of GCY receptors (Figure 4E). Although morphologically symmetric, these neurons exhibit functional asymmetry in salt-sensing responses^51,52^.

To systematically assess preservation of lineage structure in the latent space, we evaluated lineage purity and average lineage distance within randomly selected kNN neighborhoods, with k ranging from 30 to 300 (Figure 4F). We reasoned that if a latent representation reflects lineage structure, each cell’s neighbors should belong predominantly to the same lineage or an immediate relative—captured by high purity and low average lineage distance. CONCORD maintained high lineage purity even at large values of k. Furthermore, neighboring cells from different lineages were often close relatives, as reflected by a low average lineage distance. In contrast, other methods produced embeddings with significantly more mixed-lineage neighborhoods. Collectively, these findings indicate that the CONCORD latent space preserves genuine lineage structures, enabling refinement of existing annotations (Supplemental Figure 5C) and highlighting its broader utility for inferring bona fide differentiation trajectories in developmental studies^53,54^.

In addition to fate bifurcation in neuronal development, fate convergence from different lineages is a common pattern in *C. elegans* organogenesis. In the context of muscle formation, CONCORD accurately resolved how the MS, C, and D lineages converge into well-resolved sub-branches of body wall muscle, as well as rare convergence events such as the integration of ABplp/ABprp- and MS-derived cells into intestinal muscle (mu_int) (Figure 4G). Pharyngeal development—featuring complex branching and convergence of AB- and MS-derived cells—was likewise resolved in detail by CONCORD (e.g., pm3–5 deriving from both AB and MS lineages, and pm1–2, 6–8 specific to AB/MS lineages), whereas other methods recovered fewer fine-grained details (Figure 4H). Crucially, all analyses were performed directly in CONCORD’s global latent space, without subset-specific variable gene selection or re-alignment—steps that are often necessary for other methods.

Finally, to test model generalizability, we trained CONCORD and scVI on a subset of *C. elegans* batches and projected them onto unseen *C. elegans* and all *C. briggsae* data (Figure 4I). CONCORD successfully integrated the held-out batches, aligned the two species, and resolved the majority of cell types. In contrast, scVI produced a markedly lower-quality projection, with poor cross-species alignment and diminished cell type resolution.

### CONCORD captures cell cycle and differentiation trajectories in mammalian intestinal development

Unlike *C. elegans*, where early divisions are largely driven by maternal transcripts^55^, mammalian development involves extensive proliferation coupled with ongoing differentiation. To assess whether CONCORD can resolve these intertwined processes, we applied it to a single-cell atlas of embryonic mouse intestinal development^56^, which spans multiple developmental stages, batches, spatial segments, and enriched cell populations—posing a challenging integration task due to incomplete batch coverage.

CONCORD effectively integrated the data and resolved fine-grained substructures across diverse cell types (Figure 5A, Supplemental Figure 6A). Both *hcl* and *kNN* modes revealed loop-like patterns within many cell types—as evidenced by persistent homology—and often missed by other methods (Figure 5B-D, Supplemental Figure 6B). The majority of these loops correspond to cell cycle progression, supported by progressive expression of cell-cycle gene programs along the loops (Supplemental Figure 6B). For example, in intestinal epithelial cells, CONCORD not only resolved rare subtypes such as enteroendocrine cells (EECs), but also revealed two parallel trajectories—each encompassing both a cell-cycle loop and a differentiation path—corresponding to stem cell proliferation and differentiation in spatially distinct regions (Figure 5B). These structures were not captured by other methods and were supported by adult zonation markers such as *Bex4* and *Onecut2*^57^, suggesting that CONCORD can detect epithelial zonation as early as embryonic day 13.5.

**Figure 5.**
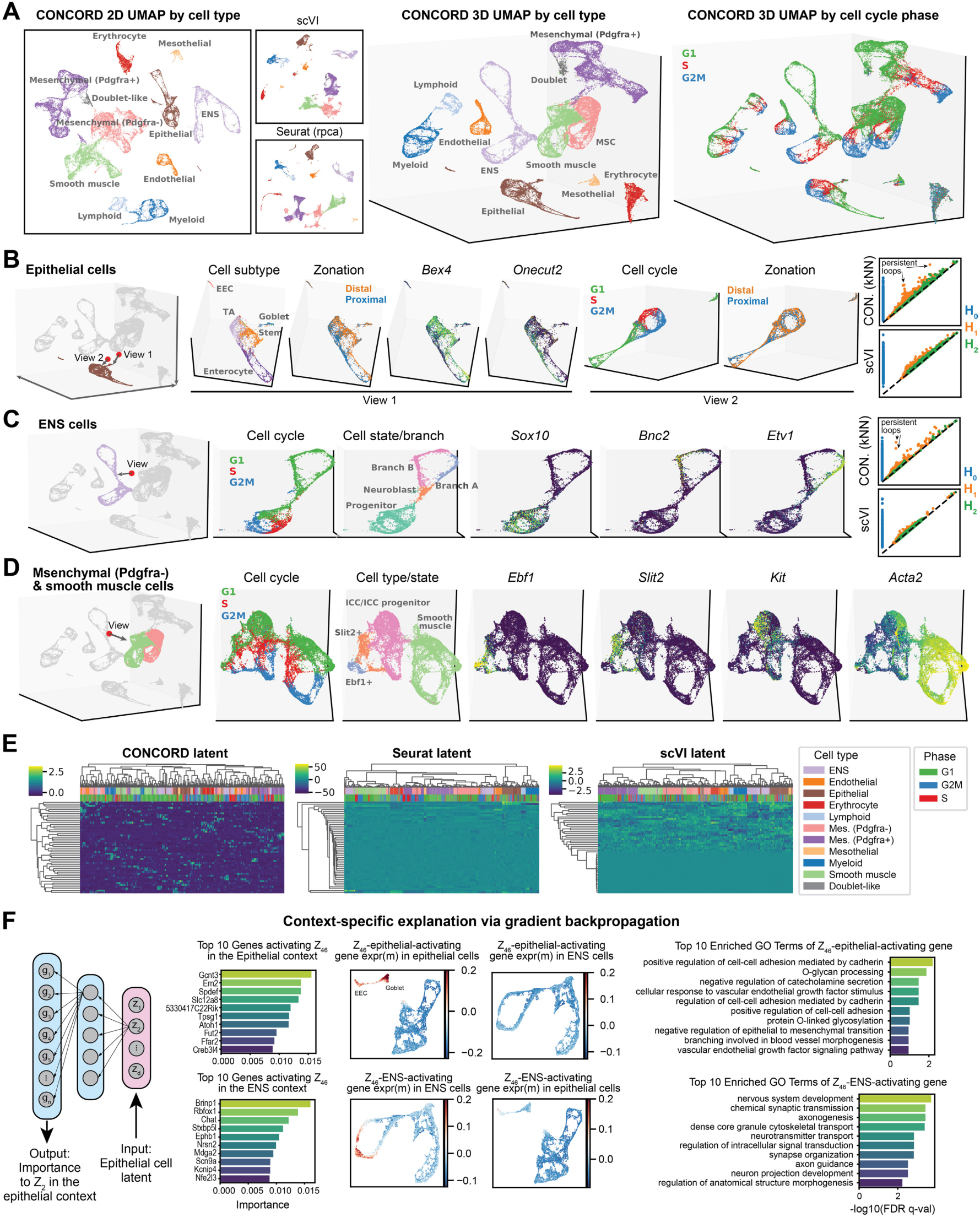
Benchmarking CONCORD on mammalian intestine development. (**A**) 2D and 3D UMAP visualizations of CONCORD (*kNN* mode) latent space, colored by cell type and cell-cycle phase, with UMAPs from scVI and Seurat (colored by cell type) for comparison. (**B**) Zoomed-in views of epithelial cells in the 3D global UMAP, colored by cell subtype, zonation, and expression of zonation-specific markers (*Bex4*, *Onecut2*). A red marker and arrow indicate the viewing angle within the 3D global UMAP. Persistence diagrams are shown for scVI and CONCORD. (**C**) Zoomed-in view of enteric nervous system (ENS) cells, colored by cell-cycle phase and cell state/branch annotations, based on Morarach et al^58^, along with state-specific marker expression. A red marker and arrow indicate the viewing angle. Persistence diagrams are shown for CONCORD and scVI. (**D**) Zoomed-in view of Pdgfra^-^ mesenchymal cells and smooth muscle cells, colored by cell-cycle phase, subtype annotation, and selected subtype-specific markers. A red marker and arrow indicate the viewing angle. (**E**) Heatmap of latent representations generated by CONCORD (*kNN*), Seurat, and scVI. (**F**) Interpretation of the CONCORD latent space using gradient-based attribution techniques. Activation of Neuron 46 (Z46) in epithelial and ENS cells is attributed to the co-expression of epithelial- and neuron-specific gene sets in their respective contexts. Gene ontology (GO) enrichment analysis of these gene sets is shown.

In the enteric nervous system (ENS), CONCORD captured the cell cycle of *Sox10*⁺ progenitor cells and identified two distinct branches of neuronal development marked by *Etv1* and *Bnc2*, matching previous observations^58^ (Figure 5C). These branches appear to converge through shared expression of neuronal maturation genes broadly active at late stages of both branches (Supplemental Figure 6C).

In mesenchymal cells—which comprise a major fraction of this dataset— CONCORD uncovered extensive heterogeneity within the *Pdgfra*⁻ and smooth muscle populations (Figure 5D). These included four consecutive cell-cycle loops marked by expression of *Ebf1*, *Slit2*, *Kit*, and *Acta2*, with gradual transitions between the loops. Notably, *Ebf1* and *Slit2* have been linked to mesenchymal multipotency^59,60^, while *Kit* marks interstitial cells of Cajal (ICC) and their progenitors^61,62^. Unlike traditional approaches where cell cycle often confounds cell type annotation, CONCORD preserves both proliferation and differentiation structure, enabling the identification of previously uncharacterized subpopulations. The complexity of these structures necessitates 3D visualization, and we encourage readers to explore the interactive embeddings: (https://qinzhu.github.io/Concord_documentation/galleries/huycke_show/).

Unlike Seurat and scVI, which left many latent dimensions underutilized, CONCORD produced a dense and interpretable latent space that reflects rich biological structure and makes full use of its representational capacity (Figure 5E). Each latent dimension typically encapsulates multiple gene co-expression programs, which can be interpreted at either single-cell or cell-state resolution using gradient-based attribution methods^63^ in a context-dependent manner (Figure 5F). For instance, latent neuron 46 (Z_46_) is activated in both epithelial cells and the ENS cells, but attribution analysis reveals it is driven by two distinct sets of highly co-expressed genes depending on the cellular context (Figure 5F, Supplemental Figure 6C). In epithelial cells, Z_46_ activation is linked to goblet cell– specific genes enriched in glycosylation pathways, whereas in ENS cells, it reflects neuronal maturation genes expressed in late-stage neurons. Notably, neither gene set shows strong expression outside its respective context, demonstrating that the CONCORD latent space captures biologically meaningful, context-specific gene co-expression programs.

### CONCORD generalizes across modalities and scales

CONCORD’s domain-agnostic design allows it to be applied to diverse data modalities beyond scRNA-seq. We tested this on a challenging single-cell ATAC-seq benchmark dataset comprising PBMCs from two donors profiled across eight different technologies^64^(Figure 6A). On both quantitative metrics and visual inspection of the embeddings, CONCORD yielded significantly better batch correction and biological label conservation than other methods, including the original study’s Harmony-based analysis (Figure 6B,C, Supplemental Figure 7A,B).

**Figure 6.**
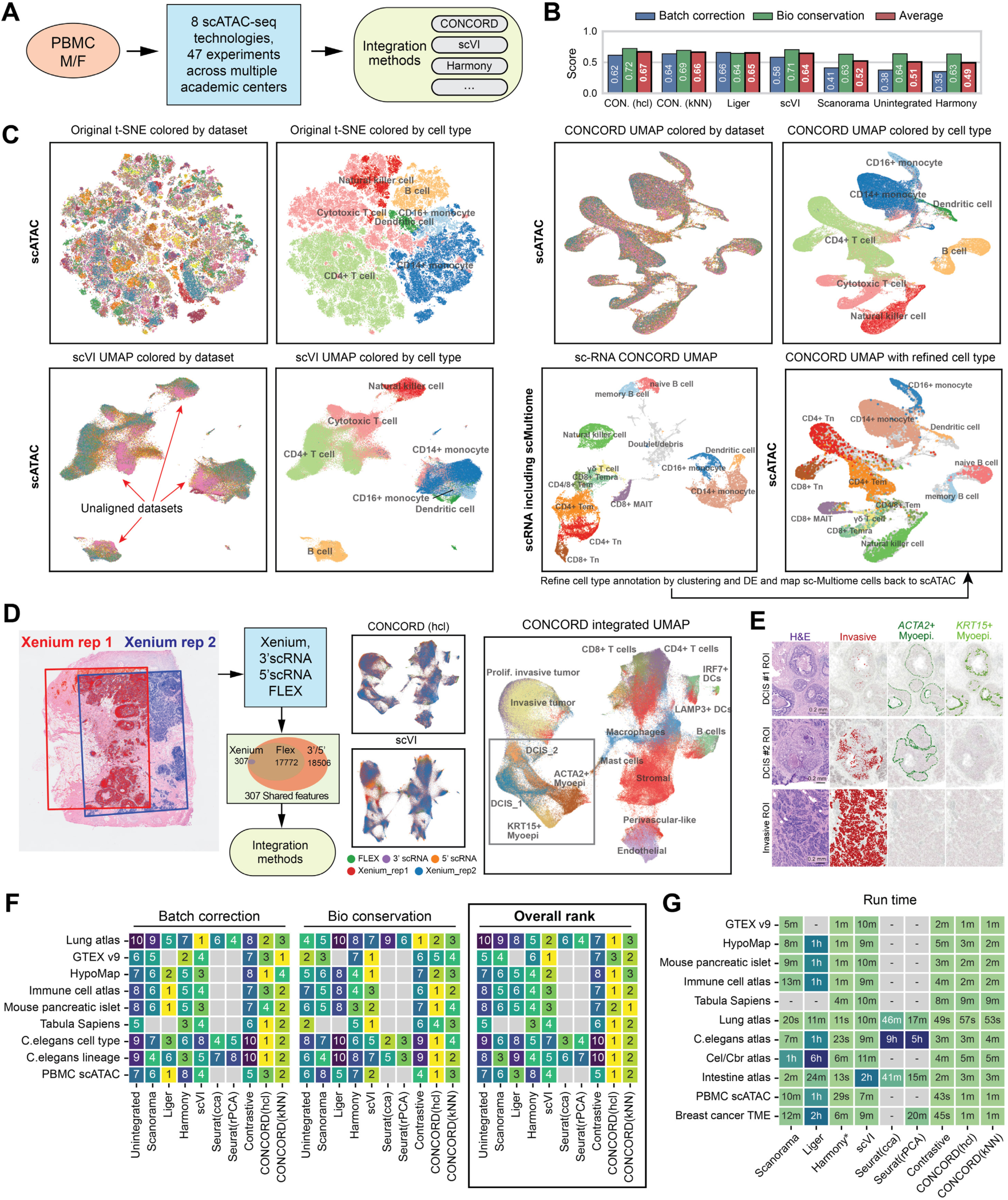
Performance of CONCORD across modalities and scales. (**A**) Schematic of the PBMC scATAC-seq benchmarking experiment spanning multiple technologies and experimental batches^64^. (**B**) Summary scores for all integration methods on the PBMC scATAC-seq data; detailed metric values are provided in Supplemental Figure 7B. (**C**) t-SNE embeddings from the original publication (Harmony integration) and embeddings produced by scVI and CONCORD, colored by batch and original cell-type annotations. To refine annotations, we analyzed paired scRNA-seq datasets with CONCORD and projected the refined labels onto the scATAC-seq embedding via shared scMultiome cells. (**D**) Schematic of the experimental design for the breast cancer tumor microenvironment sample, where a single FFPE tissue block was analyzed with multiple technologies^65^. UMAP embeddings derived from the CONCORD and scVI latent spaces are colored by batch and original cell-type annotations. Full results for all integration methods are shown in Supplemental Figure 7C. (**E**) H&E image and overlay of cell-type annotations based on Xenium data, reproduced under the Creative Commons Attribution 4.0 International License from the original publication^65^ without modification. Scale bar = 0.2 mm. (**F**) Ranking of integration method performance across all real-world benchmarking datasets, excluding datasets where scIB metrics could not be robustly computed. Each method was scored on both batch-correction and biological label conservation metrics, and the overall rank was computed based on the average score. Missing values indicate the method failed to run due to excessive resource requirements or violated assumptions. (**G**) Runtime of integration methods across all real-world benchmarking datasets. *Harmony was run using a reduced-dimensional PCA projection, whereas all other methods were applied to gene-expression matrices with 5,000–10,000 variable features.

The CONCORD embedding revealed fine-grained immune subtypes not present in the original annotations. To validate these, we refined the cell type labels using paired scRNA-seq and scMultiome data and projected them back onto the scATAC-seq embedding via shared scMultiome cells. Strikingly, refined clusters in scRNA-seq (e.g., naïve and memory B cells) corresponded precisely to clusters uncovered by CONCORD in scATAC-seq. This validation also uncovered a mis-annotation in the original study, where CD8⁺ naïve T cells were incorrectly labeled as CD4⁺ T cells. Therefore, CONCORD significantly improved analysis on existing scATAC datasets. Notably, CONCORD achieved this high-resolution result using only simple log-normalization, forgoing the complex, modality-specific data transformations often required for scATAC-seq analysis.

When applied to a breast cancer tumor–microenvironment sample profiled with Xenium, 3′ and 5′ scRNA-seq, and Fixed RNA Profiling (FLEX) technologies^65^— sharing only 307 genes—CONCORD in *hcl* mode achieved markedly better integration and cell-type resolution than other approaches (Figure 6D, Supplemental Figure 7C). A key finding of the original study was that two DCIS (ductal carcinoma in situ) subtypes exhibit distinct adjacent microenvironments: DCIS-1 is bordered by both KRT15⁺ and ACTA2⁺ myoepithelial cells, whereas DCIS-2 is encircled exclusively by ACTA2⁺ myoepithelial cells (Figure 6E). Notably, without access to spatial coordinates, CONCORD recapitulated these adjacency patterns by revealing differential connectivity between DCIS and myoepithelial clusters—consistent with signal bleed or segmentation-related transcript carryover commonly observed in spatial single-cell assays^66^.

Finally, we benchmarked CONCORD on six additional scRNA-seq datasets curated by the Open Problems in single cell analysis initiative^67^, including Tabula Sapiens (>1 million cells)^68^. CONCORD consistently achieved top performance across these datasets (Figure 6F, Supplemental Table 2) while running substantially faster and with modest RAM/VRAM requirements (Figure 6G, Supplemental Figure 7D). By contrast, several methods—including LIGER, Scanorama, and Seurat—failed to run at atlas scale owing to heavy resource demands or violations of method assumptions. CONCORD-derived 2D UMAP embeddings for these datasets are provided in Supplemental Figure 8, and additional examples, tutorials, and resources are available on the CONCORD project website (https://qinzhu.github.io/Concord_documentation/).

## Discussion

Mini-batch gradient descent underpins modern machine learning—including large language models, foundation models, and diffusion models. Growing evidence suggests that the composition of these mini-batches can influence model performance^31,69^. In contrastive learning, where each sample is contrasted against others within a mini-batch, this effect is amplified—especially in biological datasets spanning multiple batches, where naive sampling can exacerbate batch effects and distort learned representations. Yet in contrastive learning for single cell data, uniform random sampling remains the norm, limiting the method’s ability to capture biologically meaningful structure.

Our central insight is that, in contrastive learning, mini-batch composition not only influences, but fundamentally shapes the outcome. By rethinking how mini-batches are assembled, we turn contrastive learning’s sensitivity to mini-batch composition into a strength—transforming a conventional self-supervised framework into a powerful, generalizable approach for denoising, dimensionality reduction, and batch integration.

At the core of CONCORD is a unified probabilistic sampler that integrates hard-negative sampling with dataset-aware sampling. Hard-negative sampling markedly enhances the representational power of the contrastive model, enabling it to capture intricate gene co-expression programs that separate closely related cell states. The dataset-aware sampler enriches each mini-batch with cells from a single dataset, allowing the model to learn biological variation without entangling batch effects. Unlike traditional methods that rely on overlapping states or explicit batch-distortion models, CONCORD mitigates batch effects solely through principled sampling and training. As a result, it aligns cells based on shared co-varying features—a hallmark of single cell data^6,70,71^— making it especially robust when datasets have minimal overlap or unusual geometric and topological structures. The dataset-aware strategy integrates seamlessly with either the *hcl* or *kNN* hard-negative sampling variants, with both configurations yielding robust batch correction and faithful structure preservation across diverse benchmarks.

Importantly, CONCORD achieves state-of-the-art performance using a minimalistic encoder architecture—just a single hidden layer—demonstrating that substantial gains can be achieved through rational sampling and training alone, without relying on deep architectures, complex objectives, or supervision. Across both simulated and real datasets with different scale and modalities, CONCORD consistently learns latent spaces that are denoised, interpretable, and topologically faithful. In whole-organism embryogenesis atlases, it accurately reconstructs fate bifurcations and lineage convergences, enabling detailed tracing from progenitor cells to terminal states. In contrast, existing methods often misalign these datasets, lose resolution, or fragment continuous trajectories. In mammalian intestinal development, CONCORD captures complex hierarchies, spatial zonation, and cell cycle loops—all within a single integrated analysis. Unlike traditional workflows that regress out cell cycle effects, CONCORD preserves and resolves both proliferative and differentiation programs, facilitating investigations into their interplay. Its interpretable latent space further enables gradient-based attribution analyses, allowing gene-level mechanistic insights at single-cell or cell-type resolution.

CONCORD features a speed-optimized, memory-efficient design. Key components—including a vectorized sampling algorithm, native sparse matrix support, and out-of-core data loading—enable it to readily analyze million-cell atlases that may exceed available system memory. While the current implementation emphasizes simplicity, the framework is also fully extensible to more complex architectures—such as deeper neural networks or transformers^72^—to support more intricate data modalities or biological contexts. This minimalist design reduces the number of tunable parameters, though several hyperparameters remain critical for optimal performance. Although we provide default parameter settings and recommended parameter ranges based on extensive benchmarking, users may need to tune key parameters for optimal performance for untested scenarios and modalities.

Beyond the core contrastive encoder, CONCORD supports optional decoder and classifier modules for gene-level batch correction, label transfer, and annotation-guided learning. Preliminary results suggest these tasks benefit from the model’s robust latent space, though further validation is ongoing. Owing to its domain-agnostic design and generalized sampling framework, CONCORD holds potential beyond single-cell biology, offering a flexible and powerful approach for representation learning across a wide range of domains.

## Methods

### Self-supervised contrastive learning and sparse coding

We implemented CONCORD in PyTorch, building on a self-supervised contrastive learning approach inspired by SimCLR^21^ and SimCSE^22^, but with a unique dataset- and neighborhood-aware sampler design. The core objective is the normalized temperature-scaled cross entropy (NT-Xent)^21,22,73^, applied to cell representations generated via random masking, following unsupervised SimCSE^22^.

Theoretically, contrastive learning with ReLU networks and random masking augmentation can provably recover underlying sparse features from data approximated as:

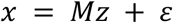

where *Mz* represents the sparse signal with ‖*z*‖_0_ = *Õ*, (1), and *ɛ* denotes noise^29^. CONCORD adopts similar conditions: we use LeakyReLU activations and independent random masking augmentations to capture correlated gene co-expression patterns while suppressing noise.

This sparse coding approach generalizes beyond traditional methods like non-negative matrix factorization (NMF), principal component analysis (PCA), factor analysis, and variational autoencoders (VAEs). Unlike these:

- It does not enforce orthogonality on *M* (as in PCA),
- It does not require non-negativity constraints (as in NMF),
- It does not assume a probabilistic generative model (as in factor analysis and VAEs),
- It does not enforce Gaussian priors on the latent space (as in standard VAEs).

Instead, it assumes an intrinsic low-rank structure shaped by gene co-expression programs, supported by single-cell studies^6,70,71^. By relaxing constraints on orthogonality, non-negativity, and Gaussian priors, the contrastive learning framework is better positioned to capture diverse gene regulatory programs that deviate from conventional assumptions. Furthermore, random masking enhances robustness to scRNA-seq dropout and improves biological interpretability, allowing the latent space to represent gene programs faithfully.

### Model architecture

CONCORD emphasizes architectural flexibility and minimalism, using a single-hidden-layer encoder for all presented benchmarking analyses to show gains from sampling and training alone. However, the architecture is fully extensible: users may substitute the encoder with more advanced models—such as deeper neural networks or transformers—to accommodate different data modalities or capture higher-order biological structures.

#### 1. Data augmentation

Input gene-expression values are normalized by total count and log-transformed to ensure consistency. We apply two complementary augmentation strategies within each mini-batch to enhance robustness and encourage reliance on redundant gene programs:

- Feature-wise masking: Randomly sets the expression of a specific gene to 0 across all cells in the mini-batch, simulating gene dropout.
- Element-wise masking: Randomly sets the expression of a specific gene to 0 for individual cells, mimicking localized noise or missing data.

Both techniques compel the encoder to leverage correlated gene co-expression patterns rather than depending on individual genes, improving generalization and noise resilience.

#### 2. Encoder

The encoder maps masked gene expression vectors to low-dimensional embeddings. By default, it is a fully connected network with one hidden layer, though the number of layers or neurons can be customized. An optional learnable feature-weighting module can precede it to weight genes sparsely and interpretably.

#### 3. Layer normalization and activation

Each linear layer is followed by layer normalization and a user-configurable activation function (default: Leaky ReLU). Layer normalization operates across features within each sample, offering robustness to batch-specific variation—making it preferable to batch normalization^74^, though the latter is also supported.

#### 4. Optional decoder and classifier

A decoder can be appended to the latent embeddings to reconstruct batch-corrected gene expression profiles. It supports joint training with the encoder or training with a frozen or partially frozen encoder. To prevent reintroduction of batch-specific variation, a distinct, learnable dataset embedding is appended during decoding, preserving the batch-effect-free nature of the core latent space.

A classification head, implemented as a multi-layer perceptron with cross-entropy loss, can be added to the encoder for tasks like cell-type annotation or doublet detection. The classifier can be trained on a pre-trained encoder or jointly with the encoder to enhance cell-type separation in the latent space. While joint training boosts class discrimination, it may impose a strong prior that disrupts trajectory continuity. To address overfitting, we recommend using a train-validation split with early stopping during classifier training.

### Contrastive objective

We adopt the noise-contrastive estimation (NCE) framework with the normalized temperature-scaled cross entropy (NT-Xent) loss^21,22,73^. Given a mini-batch of *B* cells, we generate two augmented views per sample, yielding embeddings *z_k_* and 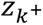 from the encoder. The standard NT-Xent loss encourages the model to pull positive pairs (different views of the same sample) closer while pushing negative pairs (views from different samples) apart. For a concatenated batch of 2*B* embeddings 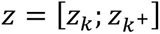, the loss is computed as:

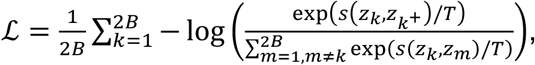

where 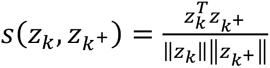 is the cosine similarity, and *T* is a customizable temperature hyperparameter that controls the trade-off between local separation and global uniformity of the embeddings^75^. The denominator sums over all other embeddings in the batch, approximating negatives sampled from the data distribution *P*. The loss is efficiently implemented using matrix operations: the logit matrix *L* = *zz^T^*/*T* is computed, diagonal entries are set to −∞ to exclude self-similarities, and cross-entropy is applied with positive indices corresponding to 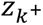 for each *z_k_*.

### Dataset and neighborhood-aware probabilistic sampler

At the heart of CONCORD is a probabilistic mini-batch sampler that determines how cells are grouped and contrasted during training. Unlike conventional contrastive learning frameworks that rely on uniform random sampling, CONCORD introduces a unified, generalizable sampling strategy that simultaneously (i) performs hard negative sampling in either *kNN* or *hcl* mode and (ii) restricts each mini-batch primarily to a single dataset. This principled design reshapes the outcome of contrastive learning, enabling the model to produce a coherent, high-resolution, and batch-effect-mitigated representation of the cell state landscape.

#### 1. *kNN* mode

We begin by coarsely approximating the global data manifold using a k-nearest neighbors (kNN) graph, where k is a user-defined parameter (typically moderately large). The graph can be initialized from normalized gene expression values, a PCA projection, or a CONCORD batch-corrected embedding generated with the dataset-aware sampler. By default, we run CONCORD with the dataset-aware sampler for 2 epochs, followed by the remaining epochs with joint sampling. For scalability, we leverage the *Faiss* library^76^ for efficient neighbor retrieval in large datasets. This kNN graph then guides neighborhood-aware sampling, modulated by a user-defined neighborhood enrichment probability *P_kNN_*. To construct mini-batches that are both dataset- and neighborhood-enriched, we partition each mini-batch into four subsets—in-dataset neighbors, in-dataset global samples, out-of-dataset neighbors, and out-of-dataset global samples (Figure 1F). A “core sample” is randomly selected from one dataset to anchor both neighborhood and dataset-aware sampling. The four subsets are then sampled based on *P_d_* (the probability of sampling from the same dataset) and *P_kNN_* as follows:

- In-dataset neighbors: *P_d_* ⋅ *P_kNN_* ⋅ *B* cells from the same dataset and within the core cell’s kNN neighborhood.
- In-dataset global samples: *P_d_* ⋅ (1 − *P_kNN_*) ⋅ *B* uniformly sampled cells from the same dataset, outside the neighborhood.
- Out-of-dataset neighbors: (1 – *P_d_*) ⋅ *P_kNN_* ⋅ *B* cells from other datasets that fall within the core cell’s kNN neighborhood.
- Out-of-dataset global samples: (1 – *P_d_*) ⋅ (1 − *P_kNN_*) ⋅ *B* uniformly sampled cells from all other datasets.

#### 2. *hcl* mode

Unlike *kNN* mode, which explicitly samples cells from within the kNN neighborhood, *hcl* mode implements the hard negative sampling algorithm from Robinson et al.^32^, utilizing importance sampling to compute the expected hard-negative loss directly within the contrastive objective without altering the mini-batch sampling procedure. We briefly explain the method here and refer readers to the original publication for details.

Given an anchor embedding *z_k_*, to enrich negative samples *z_m_*, we propose sampling from the mixed hard negative distribution *q_β_*(*z_m_*) ∝ exp(*βs*(*z_k_*, *z_m_*)/*T*) ⋅ *P*(*z_m_*), where the similarity 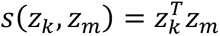 (with embeddings ℓ_2_-normalized), and *β* > 0 is a “concentration parameter”. The exponential term acts as a von Mises–Fisher kernel: larger *β* concentrates mass on points whose embeddings already lie close to the anchor (harder negatives), while *β* = 0 recovers the uniform sampler over the data distribution *P*.

Since sampling directly from *q_β_*is computationally inefficient, we apply importance weights in the contrastive loss to approximate the expected contribution under *q_β_*. Specifically, the contrastive loss under *hcl* mode is:

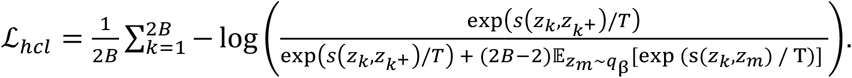

Note that this omits an optional debiasing term (from Equation 4 in Robinson et al.^32^), which subtracts potential contamination from false positives (same-class negatives) using the class prior *τ*_+_. We set *τ*_+_ = 0, as single-cell data are highly heterogeneous, making it unlikely to sample cells with identical molecular states within a mini-batch.

The expectation is computed using Monte-Carlo importance sampling:

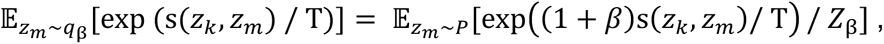

where 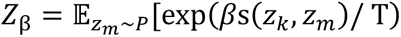 is the partition function, empirically estimated as:

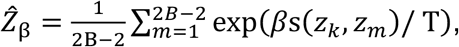

Thus,

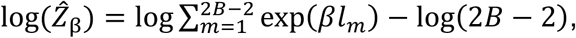

with *l_m_* = *s*(*z_k_*, *z_m_*)/*T* as the original negative logits. This leads to reweighted negative logits 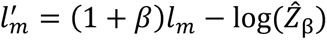, which replace the originals in the NT-Xent denominator.

To integrate the *hcl* hard-negative sampler with the dataset-aware sampler, apply the *hcl* contrastive loss to mini-batches constructed under the dataset-aware probability distribution (determined by *P_d_*) instead of the uniform data distribution *P*, enabling simultaneous batch correction by focusing contrasts primarily within datasets. In practice, we find that the *hcl* mode is more sensitive to the dataset-aware sampling probability (*P_d_*), performing optimally with strict intra-dataset sampling (*P_d_* = 1.0). We hypothesize this is due to *hcl*’s strong emphasis on close neighbors: when such neighbors are drawn across batches, they may represent the same biological state, leading the model to inadvertently penalize correct alignments.

Both variants of CONCORD samplers are implemented using vectorized operations in PyTorch and NumPy to optimize memory efficiency and minimize computational overhead, ensuring scalability across large datasets and enabling rapid training.

### Model training

Mini-batches are constructed via the sampler, shuffled, and optimized with NT-Xent using the Adam optimizer^77^. Interestingly, and in contrast to trends often seen in computer vision, we found that CONCORD’s performance did not improve with very large batch sizes (relative to the total number of cells). Instead, for the *C.elegans* dataset (>90,000 cells), performance peaked at moderate sizes (e.g., 256-512) and then began to decline as the batch size exceeded 1,000 (Supplemental Fig. 4E). We hypothesize that this is because the benefit of hard-negative sampling strategy is diluted in very large batches. For resolving the fine-grained distinctions common in biological data, the model requires an enrichment of “hard negative” contrasts. As batch size increases, the mini-batch distribution approaches the global data distribution, reducing the effectiveness of this targeted sampling. Therefore, for all benchmarking analyses, we adopted a moderate default batch size of 256, which not only achieves optimal performance across our diverse benchmarks but also makes CONCORD highly accessible. This choice ensures that VRAM requirements are minimal (Supplemental Fig. 7D), allowing the method to be run efficiently on widely available, cost-effective GPUs.

Besides the core contrastive objective, optional loss terms, including mean squared error (MSE) for the decoder, cross-entropy loss for classification, and L1/L2 regularization for feature-weighting modules, can be incorporated with adjustable weights. A learning rate scheduler is employed to gradually reduce the learning rate over time, promoting stable convergence.

### Simulation pipeline

We developed a versatile simulation pipeline to generate synthetic single-cell gene expression data with diverse underlying structures. Unlike conventional simulators that predominantly produce discrete clusters, our pipeline accommodates a broad range of topologies, including linear trajectories, branching trees, loops, and intersecting paths frequently observed in real single-cell datasets.

The pipeline proceeds in three sequential stages, as illustrated in Figure 2A.

#### 1. Groundtruth data model

In the first stage, the state simulator constructs a noise-free data matrix [N × D] (where N is the number of cells and D is the number of genes) according to a user-defined structure:

- Clusters: Cells form discrete groups characterized by unique gene programs, optionally including shared or ubiquitously expressed genes.
- Trajectories: Cells exhibit gradual shifts in gene expression, emulating cell differentiation processes.
- Loops and intersecting paths: Continuous trajectories that close into loops or intersect, representing cyclic biological processes.
- Trees: Hierarchical, branching lineages representing progenitor-to-terminal fate differentiation, configurable by branching factor and tree depth.

#### 2. Noise model

Expression values are then sampled from selected distributions (e.g., Gaussian, Poisson, Log-normal, Negative Binomial), introducing realistic variability and dropout patterns. Users can precisely control parameters including mean baseline expression, dispersion (noise level), dropout probability, and can enforce non-negativity or integer rounding of the generated values, yielding a noisy data matrix [N × D].

#### 3. Batch model

In the final stage, an optional batch simulator introduces dataset-specific technical variability to mimic batch effects. For each batch, a user-specified effect type is applied to a subset of cells (determined by customizable proportions), enabling simulation of various technical artifacts. Supported effect types include:

- Variance inflation: Multiplies each entry by 1 + *N*(0, *σ*^2^), where *σ* is the dispersion parameter.
- Batch-specific distribution: Adds noise sampled from a specified distribution (e.g., Normal, Poisson, Negative Binomial, Log-normal) with configurable mean and dispersion.
- Uniform dropout: Randomly sets a fixed fraction of values to zero.
- Value-dependent dropout: Drops values with probability 1 − exp(−λ ⋅ *x*^2^), where *λ* is the level parameter and *x* is the expression value.
- Down-sampling: Subsamples UMI counts to a specified ratio, simulating reduced sequencing depth.
- Scaling factor: Multiplies the entire matrix by a scalar to shift overall expression levels.
- Batch-specific expression: Adds distribution-based noise to a random subset of genes.
- Batch-specific features: Appends new genes unique to the batch, with expression sampled from a specified distribution.

Multiple simulated batches are then concatenated into a single dataset, with varying degrees of batch overlap to mimic real-world sampling scenarios, producing the final input data matrix [N × D] with noise and batch effects.

By combining diverse gene expression structures with realistic noise models and customizable batch effects, this simulation pipeline can approximate a broad spectrum of biological and technical scenarios. As such, it provides a powerful testbed for benchmarking data-integration techniques, trajectory-inference algorithms, and manifold-learning methods under controlled yet biologically realistic conditions.

### Benchmarking pipeline

To comprehensively evaluate the performance of CONCORD and other dimensionality-reduction or data-integration methods, we designed a robust benchmarking pipeline that integrates geometric, topological, biological label conservation, and batch-correction metrics. This multifaceted assessment framework consists of the following components:

#### 1. Topological assessments

To quantify the preservation of intrinsic topological features, we employed persistent homology analysis implemented via Giotto-TDA^78^. Persistent homology captures structural properties of the data across multiple scales, using Vietoris-Rips complexes constructed over increasing radii to generate persistence diagrams and Betti curves. Persistence diagrams reveal the lifespan of topological features such as connected components (Betti-0), loops (Betti-1), and voids (Betti-2). We derived Betti curves from these diagrams, interpolating them onto a common filtration grid (number of bins = 100) for consistency. For each homology dimension, we computed the mode of the Betti curve (representing the most persistent Betti number across scales) and compared it to known topological ground truths using L1 distance across dimensions, yielding a Betti number accuracy score of 1/(1 + L1). Additionally, we computed the stability of Betti curves as 1/(1 + variance), where variance is the variance of the Betti values across the filtration grid. The Betti curve stability score is averaged across dimensions to quantify topological robustness (scores range from 0 for highly variable curves to 1 for perfectly stable ones). The final topology score is a weighted average: 0.8 × Betti number accuracy + 0.2 × Betti curve stability.

#### 2. Geometric assessments

We evaluated the preservation of geometric relationships by calculating Pearson correlations between pairwise distances in the embeddings and those in the corresponding noise-free reference data, providing a measure of global geometric fidelity. For local neighborhood preservation, we employed trustworthiness^79^, a metric that assesses how faithfully high-dimensional neighborhood structures are maintained in lower-dimensional embeddings. Trustworthiness scores range from 0 (poor preservation) to 1 (perfect preservation), and we computed average trustworthiness scores across neighborhood sizes (k-values) from 10 to 100 in increments of 10. Additionally, we visualized trustworthiness as a function of k to reveal how each method performs at different local scales.

#### 3. Batch correction metrics

We adopt established metrics from the scIB-metrics package^33^ to systematically evaluate batch correction, including:

- Graph connectivity: Assesses whether cells of the same biological label form a connected component in the integrated k-nearest neighbor (kNN) graph, with scores ranging from 0 to 1 (1 indicating full connectivity).
- iLISI (integration Local Inverse Simpson’s Index): Measures batch mixing by estimating the effective number of batches in local neighborhoods, rescaled to 0–1 (higher scores indicate better integration).
- KBET (k-nearest neighbor Batch Effect Test): Determines whether the batch label composition in a cell’s k-nearest neighborhood is similar to the expected global composition, computing a rejection rate (higher rejection indicates poorer mixing); the rate is averaged across labels and subtracted from 1 to rescale to 0–1, with higher scores denoting better batch removal.
- PCR comparison (Principal Component Regression): Quantifies the variance contribution of batch effects via regression on principal components, comparing pre- and post-integration, rescaled to 0–1.
- Silhouette batch (Batch ASW): Computes the average silhouette width (ASW) using the absolute silhouette width on batch labels per cell, subtracted from 1 (so 1 indicates perfect mixing), averaged within each cell type and then across types; ranging from 0 (strong separation) to 1 (ideal integration).

#### 4. Biological label conservation

These metrics, also from scIB-metrics^33^, evaluate the preservation of biological variance and cell-type separation:

- Isolated labels: Assesses handling of rare or batch-specific labels using F1 score and ASW, scaled to 0-1 (higher scores indicate better separation of isolated labels).
- Leiden ARI (Adjusted Rand Index): Measures overlap between true biological labels and clusters obtained via Leiden clustering on the integrated data, with scores from 0 (random) to 1 (perfect match).
- Leiden NMI (Normalized Mutual Information): Quantifies information shared between true labels and Leiden clusters, ranging from 0 (no overlap) to 1 (perfect correspondence).
- Silhouette label (Cell Type ASW): Evaluates cell-type separation using average silhouette width on labels, scaled to 0-1 (1 indicating dense, well-separated clusters).
- cLISI (cell-type Local Inverse Simpson’s Index): Estimates cell-type purity in local neighborhoods, rescaled to 0-1 (higher scores indicate better separation).

A key limitation of the scIB pipeline is that the pipeline does not fully accommodate the hierarchical and continuous nature of many biological systems. Consequently, for simulations with continuous trajectories or loops, we first apply Leiden clustering to noise-free data to define “clusters” as ground truth, or use “branch” labels as a proxy for cell states in tree simulations. Under these conditions, the scIB metrics are applied in a more coarse-grained manner, offering an approximate assessment in these more complex scenarios.

#### 5. Probing classifiers

To further assess embedding quality, we implemented probing classifiers—often used in evaluating representation learning methods. Two types of probing classifiers are implemented: a k-nearest neighbors (KNN) probe and a linear probe. The KNN probe fits a k-nearest neighbors classifier on 80% of the data and evaluates on the held-out 20%. The linear probe fits a single fully connected layer to the fixed embeddings using AdamW optimization, with cross-entropy loss for classification or MSE for regression. It uses a train-validation split (80/20) and performs early stopping based on validation loss (default patience 5 epochs) to prevent overfitting.

We use these probes for evaluating biological label conservation or batch mixing via classification. For the former task, the classification accuracy is used, while for the latter we use the classification error to evaluate the degree of batch mixing. Note that on datasets with imbalanced batch composition and coverage, high classification error could sometimes indicate over-correction. We therefore only use batch classification error to quantify batch mixing when scIB metrics failed to run (e.g., *C. elegans*/*C. briggsae* atlas), and include label classification accuracy under the biological label conservation metrics for all evaluations.

For all benchmarking analyses, we applied standard total count normalization and log transformation to the datasets, followed by selection of highly variable features (5,000 for all Open Problems datasets and 10,000 for the *C. elegans*/*C. briggsae* datasets, intestine atlas, and PBMC scATAC-seq data). The resulting normalized matrices served as input for all integration algorithms except Harmony, which requires PCA-projected coordinates. All methods were benchmarked in latent spaces of identical dimensionality: 30 dimensions for simulated datasets, 50 dimensions for most real-world datasets, and 300 dimensions for complex datasets—such as the *C. elegans*/*C. briggsae* and Tabula Sapiens atlases—to adequately capture the extensive diversity of cell states. All methods were executed on an Amazon EC2 environment equipped with an NVIDIA Tesla T4 GPU.

### Transcriptomic profiling of early *C. elegans* embryos by scRNA-seq

Wildtype N2 worms were grown on NGM plates and synchronized by bleaching. Eggs hatched on 10-cm plates and were grown until L3/L4 stage. To enrich for early embryos, plates were transferred to a 12⁰C and incubated for 48 h. Adult worms were lysed by bleaching, and embryos were dissociated into single cells as previously described^80^. Cells were loaded onto a Chromium GEM-X Single Cell 3’ Chip Kit v4 with GEM-X Universal 3’ Gene Expression v4 reagents (10x Genomics, 1000686). Libraries were prepared following the 10x Genomics protocol, sequenced on NovaSeq X, and processed with CellRanger (9.0.1) utilizing the WBcel235 transcriptome. A total of 12,899 cells were recovered, with median of approximately 69,000 reads per cell.

## Data Availability

Single-cell RNA-seq data of *C. elegans* early embryos have been deposited in the Gene Expression Omnibus (GEO) under accession number GSE305031. Public datasets analyzed in this study include: the human lung atlas, compiled by Luecken et al.^33^ and obtained from the scIB-metrics website (https://scib-metrics.readthedocs.io/en/stable/notebooks/lung_example.html); GTEX v9, HypoMap, immune cell atlas, mouse pancreatic islet, Tabula Sapiens datasets, sourced from the Open Problems in Single-Cell Analysis website (https://openproblems.bio/benchmarks/batch_integration?version=v2.0.0); the *C. elegans* embryogenesis atlas, downloaded from GEO under accession GSE126954; the joint *C. elegans* and *C. briggsae* dataset, available under GEO accession GSE292756; and the mouse intestinal developmental atlas, acquired from GEO under accession GSE233407.

## Code Availability

Concord is available at https://github.com/Gartner-Lab/Concord under the MIT License. All benchmarking codes to generate results in this manuscript are deposited to https://github.com/Gartner-Lab/Concord_benchmark. Full documentation of Concord can be found at: https://qinzhu.github.io/Concord_documentation/.

## Acknowledgement

We thank the reviewers for their constructive feedback, which was instrumental in driving major enhancements to CONCORD and strengthening the manuscript. We thank Junhyong Kim, John Murray, and Honesty Kim for valuable feedback on the manuscript. We also thank the authors of Large et al.^50^ for sharing the *C. elegans* and *C. briggsae* dataset, with special acknowledgement to Christopher R. L. Large for facilitating data access. We are grateful to authors of De Rop et al.^64^ for providing key metadata for the benchmarking analysis. Additionally, we thank members of the Gartner Lab for critical discussions, support and assistance in testing early versions of CONCORD. We acknowledge the use of ChatGPT and Gemini for assistance with code refinement and annotation.

This work was supported by grants from the NIH (U01CA199315, R01GM135462, R01DK126376, U01DK103147, and R33CA247744), the Chan Zuckerberg Initiative (CZI 2023-332284), and the UCSF Center for Cellular Construction (DBI-1548297), an NSF Science and Technology Center. Q.Z. is supported by a Cancer Research Institute Immuno-Informatics Postdoctoral Fellowship (CRI5054). Z.J.G. is a Chan Zuckerberg BioHub San Francisco Investigator. Sequencing was performed at the UCSF Center for Advanced Technology (CAT), supported by UCSF PBBR, RRP IMIA, and NIH 1S10OD028511-01 grants.

## Author Contributions

Q.Z. and Z.J.G. conceived the project. Q.Z. designed and implemented the method. Q.Z. and Z.J conducted benchmarking analyses. B.Z. collected the new *C. elegans* dataset under the supervision of L.W and Z.J.G. M.T. provided critical feedback and methods for topological data analysis. Z.J.G. supervised the project. Q.Z. and Z.J.G. wrote the manuscript. All authors reviewed and edited the manuscript.

## Supplementary Information

### Supplementary Figures

**Supplemental Figure 1.**
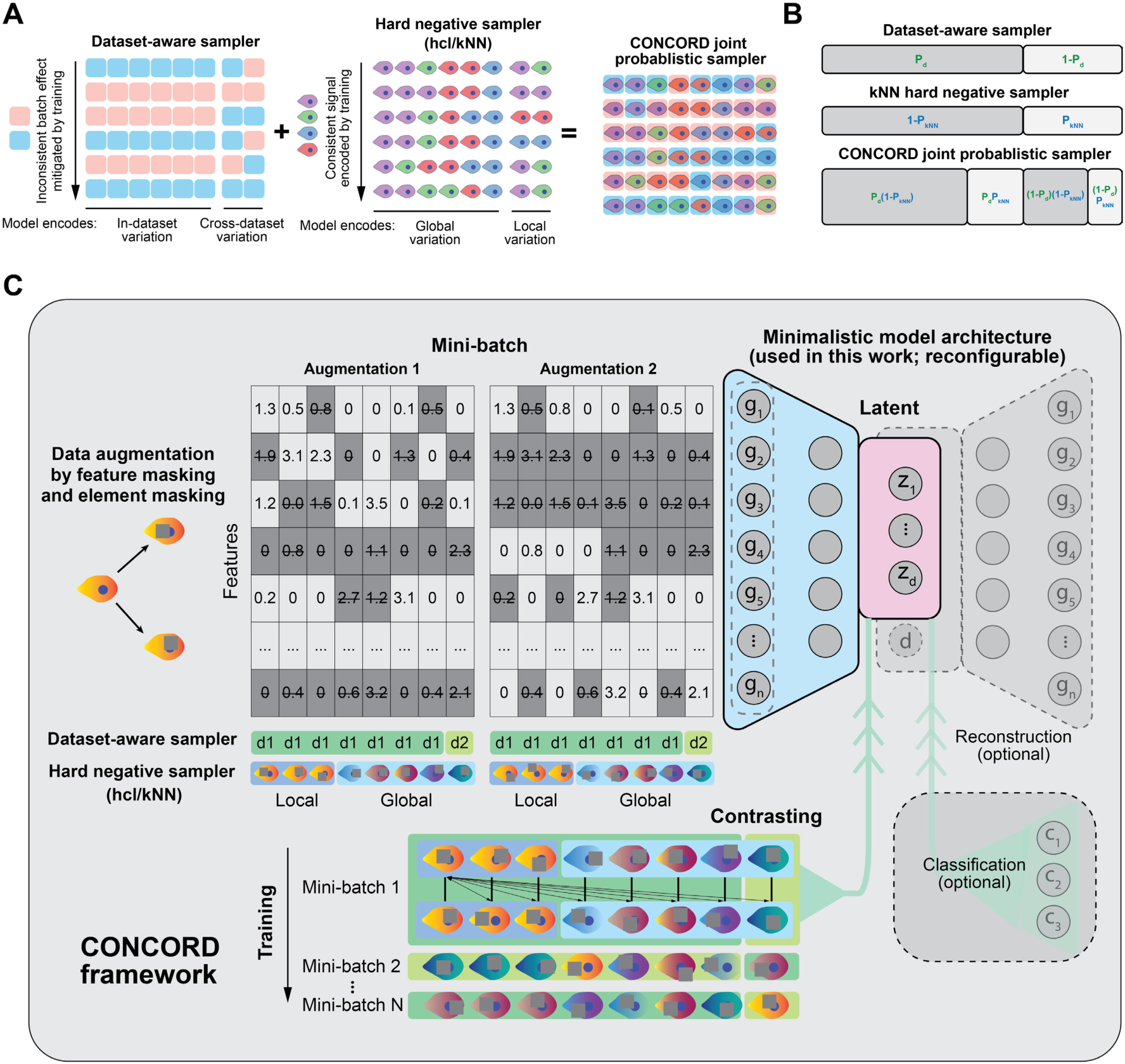
The CONCORD framework. (A) The CONCORD mini-batch sampling framework. A dataset-aware sampler that enriches each mini-batch with within-dataset cells is combined with a hard-negative sampler to support both data integration and enhanced resolution. Two hard-negative variants can be used with the dataset-aware sampler: the *kNN* sampler performs explicit local and global sampling via a kNN graph of cells; the *hcl* mode computes the expected hard-negative loss directly within the contrastive objective (see Methods). (B) Joint probabilistic sampler (kNN mode). We construct a probabilistic mini-batch sampler in which the likelihood of selecting a cell reflects the combined probabilities of dataset-aware (P_d_) and neighborhood-aware sampling (P_kNN_). (C) During training, each cell is augmented twice via random feature-wise and element-wise masking, and the contrastive loss is computed on latent encodings of mini-batches drawn by the joint sampler. The model architecture used here is a minimalist neural network with a single hidden layer, though the framework supports more complex designs. Optional modules include a decoder with a learnable dataset/batch covariate for batch-free gene-expression inference and a classifier for cell-type classification or annotation-guided learning.

**Supplemental Figure 2.**
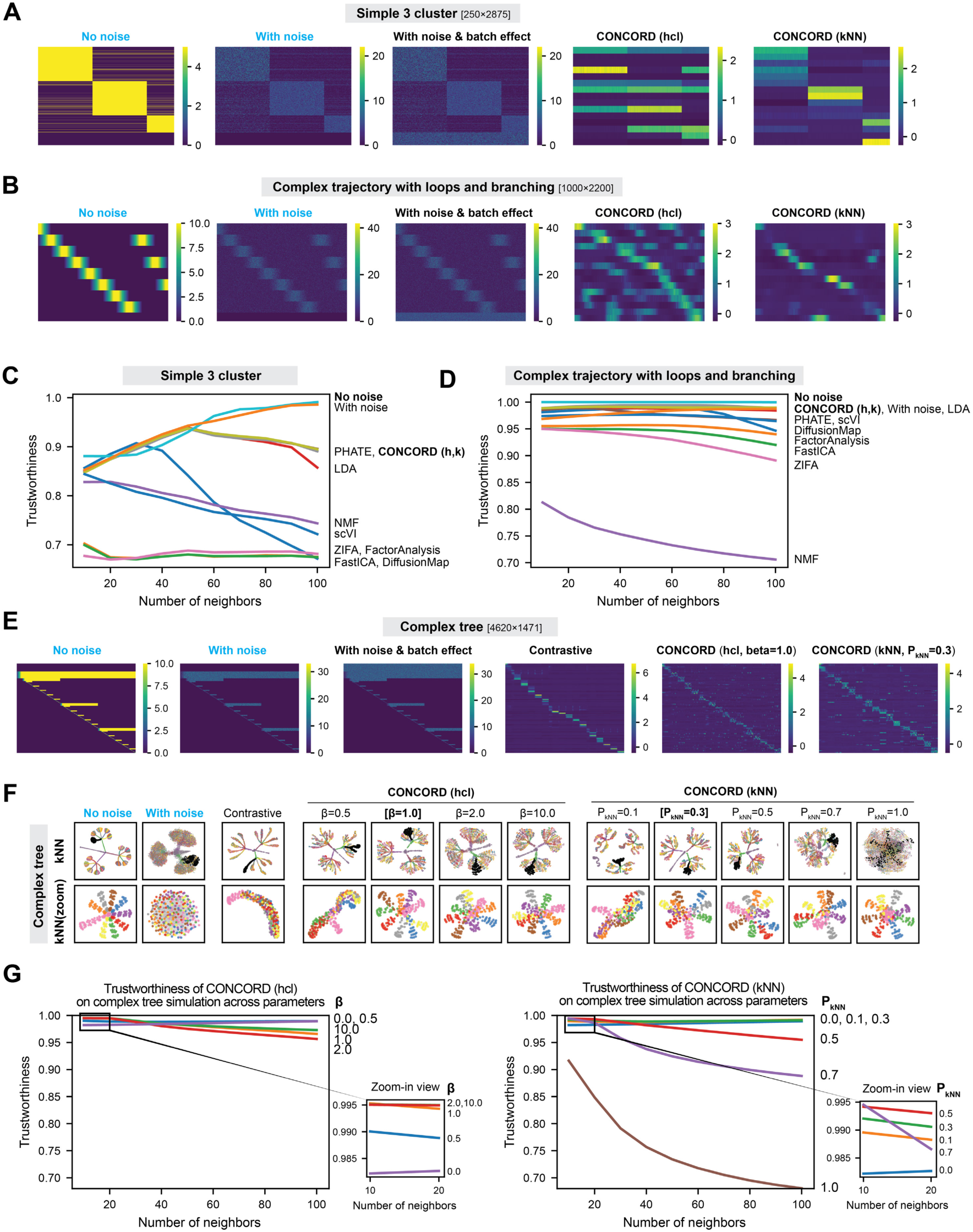
Benchmarking CONCORD and other dimensionality reduction methods across diverse structures. **(A)** Heatmaps of simulated expression for the three-cluster structure and the corresponding CONCORD latent encoding in *hcl* or *kNN* modes. (**B**) Heatmaps of simulated expression for the trajectory-loop structure and the corresponding CONCORD latent encoding in *hcl* or *kNN* modes. (**C**) Trustworthiness measured across neighborhood sizes (*k*) in the three-cluster simulation. In the noise-free reference, within-cluster neighbors are assigned at random, so trustworthiness is < 1. CONCORD (h, k) denotes the *hcl* and *kNN* modes, respectively. (**D**) Trustworthiness measured across neighborhood sizes in the complex trajectory-loop simulation. (**E**) Heatmaps of simulated expression for the complex-tree structure shown in Figure 2G and the CONCORD latent encoding under medium hard-negative enrichment in *hcl* and *kNN* modes. (**F**) Trustworthiness across neighborhood sizes for *hcl* and *kNN* modes in the complex-tree simulation, evaluated under varying degrees of hard-negative sampling. An inset for *k* < 20 highlights improved local neighborhood preservation with hard-negative sampling.

**Supplemental Figure 3.**
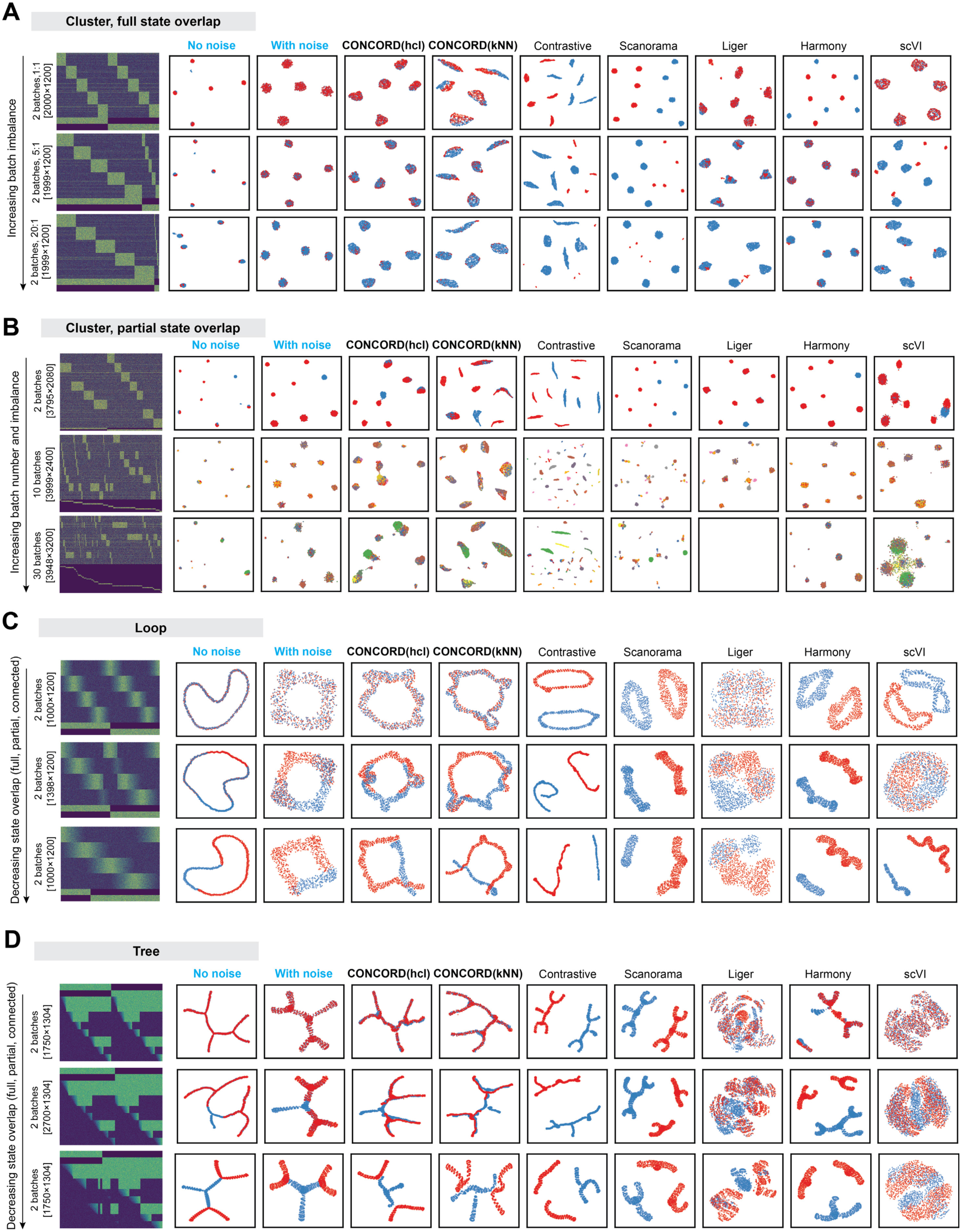
Benchmarking CONCORD and other data-integration methods across diverse structures. (**A**) Two-batch, five-cluster simulations with increasing batch-size imbalance and full state overlap. Heatmaps of the input data (dimensions indicated) and UMAPs of the ground truth and each method’s latent space, colored by batch, are shown. (**B**) Cluster simulations with increased batch number and imbalance, with partial overlap of cell states across batches. Heatmaps of inputs and UMAPs of the ground truth and each method’s latent space, colored by batch, are shown. (**D**) Loop simulations with varying batch overlap. kNN graphs (k = 15; edges omitted) colored by batch are shown for the ground truth and each method. (**E**) Tree simulations with varying batch overlap. kNN graphs (k = 30; edges omitted) colored by batch are shown for the ground truth and each method.

**Supplemental Figure 4.**
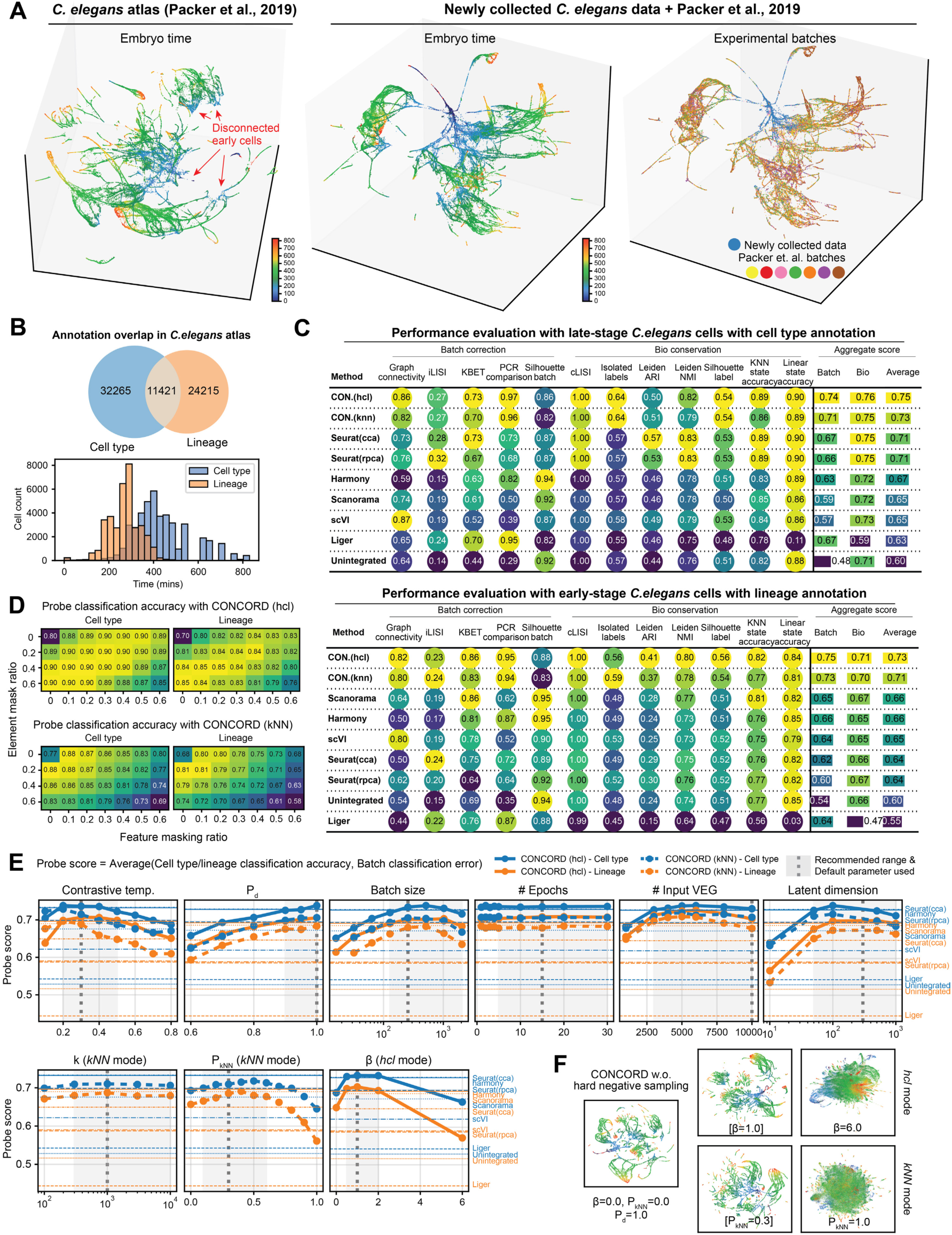
Performance of CONCORD on *C. elegans* atlas. (**A**) UMAPs of the *C. elegans* atlas from Packer et al. generated from the CONCORD latent space. Gaps among early-stage cells are apparent; adding our newly collected *C. elegans* dataset enriched for early embryos fills these gaps, yielding continuous trajectories. The combined UMAP is colored by inferred embryo time and batch. (**B**) Overlap between expert-curated cell-type and lineage annotations. A histogram shows that lineage annotations are concentrated in early-stage cells, whereas cell-type annotations are predominantly in late-stage cells. (**C**) Integration performance of CONCORD and other methods, evaluated separately for early-stage cells with lineage annotations and late-stage cells with cell-type annotations. See Methods for details on metric definitions. (**D**) Performance of the two CONCORD modes (*hcl* and *kNN*) across combinations of element- and feature-masking ratios, assessed by average classification accuracy using linear and kNN probes. (**E**) Performance of both modes across key hyperparameters, quantified by the average of label-classification accuracy and batch-classification error. Each run varies one hyperparameter while fixing the rest (default value indicated on plot). Scores from other methods are included for comparison. (**F**) UMAPs illustrating results with the dataset-aware sampler alone (no hard negatives) and with moderate versus excessive hard-negative sampling for *hcl* and *kNN* modes. Moderate local sampling improves cell-type and lineage resolution, whereas excessive local sampling without balanced global sampling disrupts global structure.

**Supplemental Figure 5.**
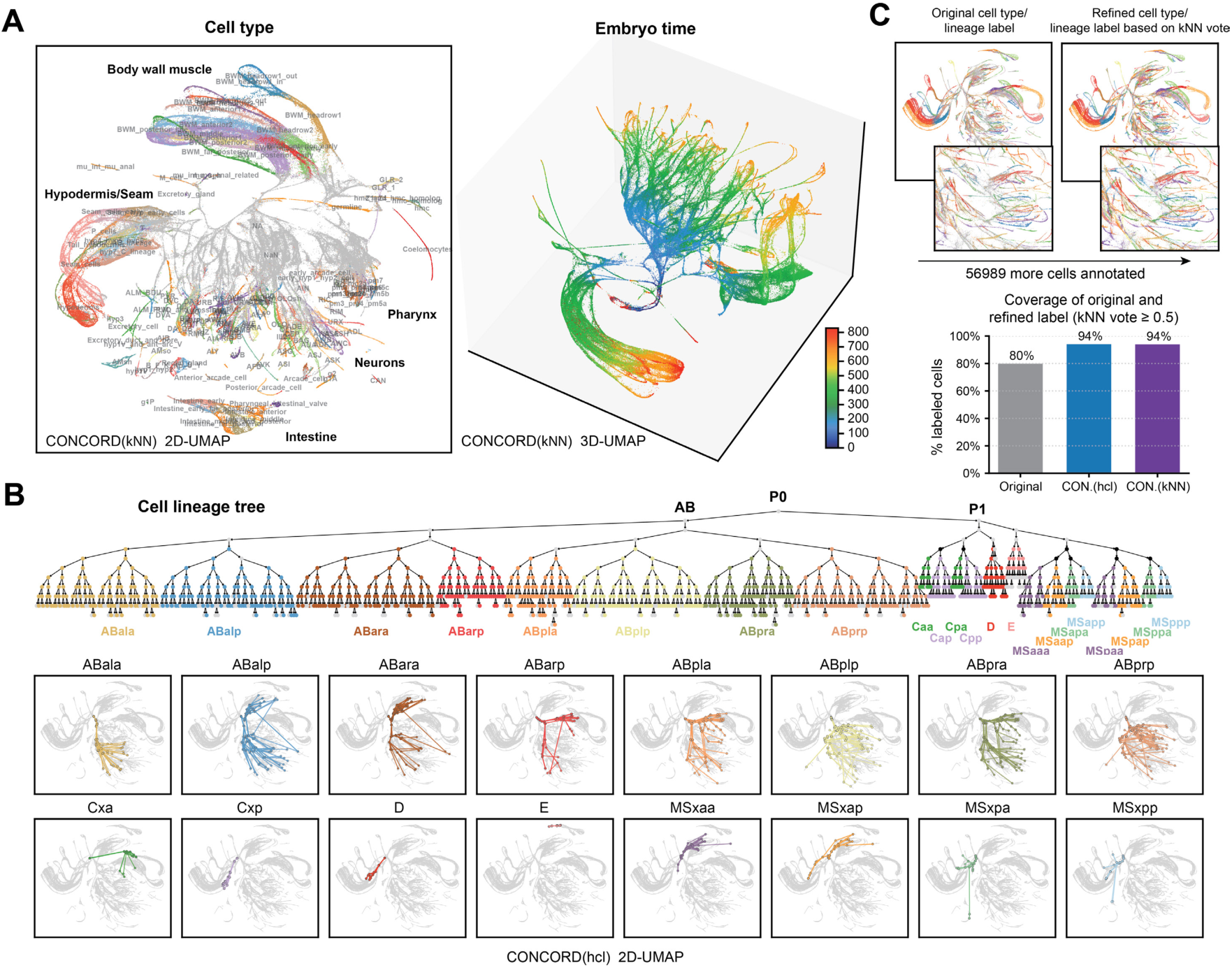
CONCORD analysis of *C. elegans/C.briggsae* embryogenesis atlas. (**A**) 2D and 3D UMAPs of the CONCORD latent space (kNN mode), colored by cell type and inferred embryo time. (**B**) *C. elegans* lineage tree and its projection onto the CONCORD (hcl) embedding. Lineage annotations from Large et al. were mapped to the *C. elegans* lineage tree (with some ambiguous mappings due to symmetry). Each lineage is represented by its cluster medoid on the UMAP; lines connect each parent lineage to its daughters following the lineage tree. Subtrees for major lineage groups are shown separately. (**C**) Label refinement in the CONCORD latent space via kNN majority vote. For each cell, we examine its k = 30 nearest neighbors; if ≥50% of neighbors carry expert-curated lineage/cell-type labels, we assign the neighborhood’s majority label to unlabeled cells (and relabel when the majority disagrees). We iterate this procedure twice so newly assigned labels can vote. This recovers labels for many unlabeled cells and flags likely mis-annotations.

**Supplemental Figure 6.**
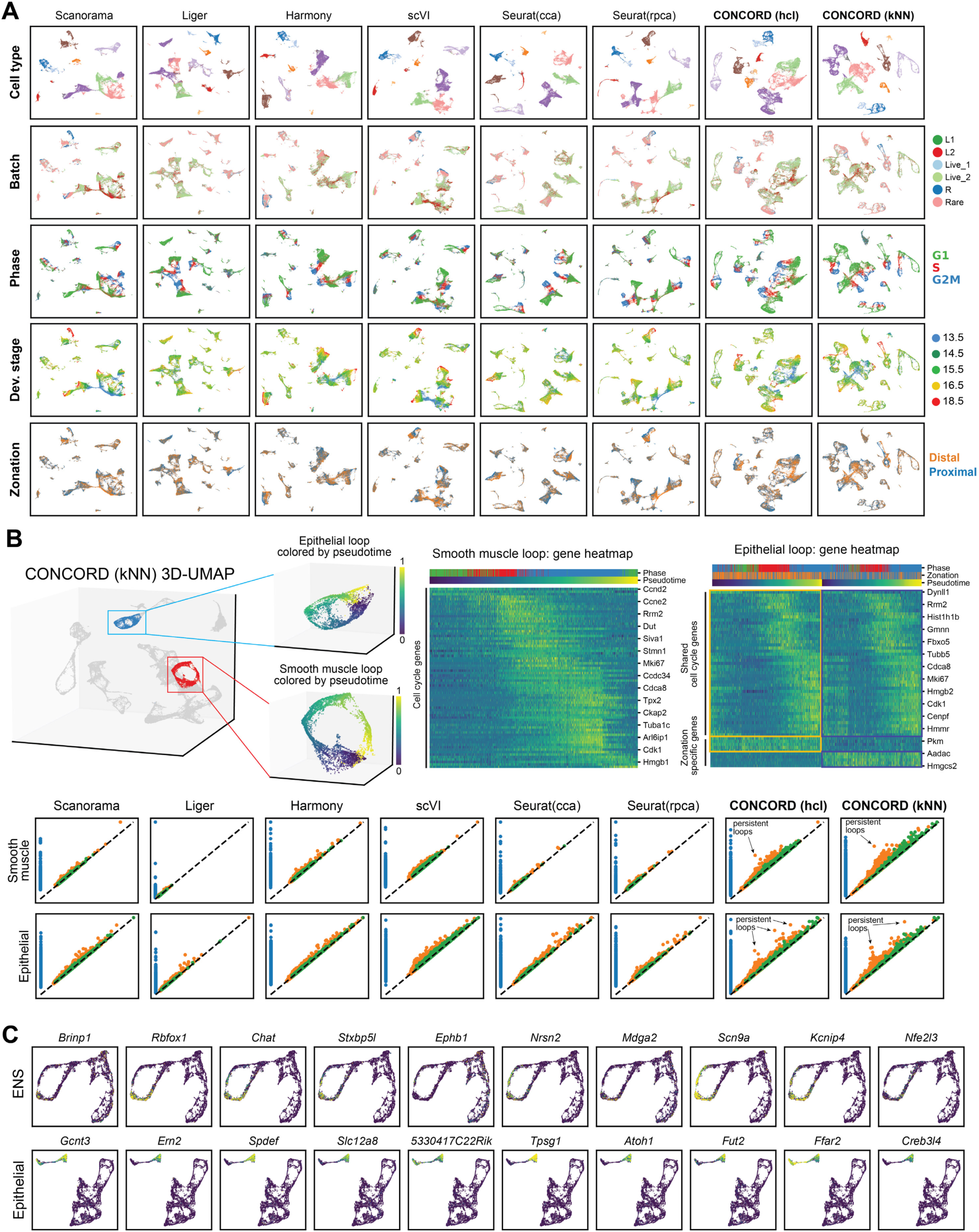
Benchmarking CONCORD on mammalian intestine development. (**A**) UMAP embeddings derived from the latent spaces of CONCORD and other integration methods for the mouse intestinal developmental atlas^56^, colored by broad cell type, batch, cell-cycle phase, developmental stage, and zonation. (**B**) For epithelial and smooth-muscle cells, loop-like trajectories were identified and pseudotime was assigned along each circular path. Heatmaps show top differentially expressed genes (DEGs) along each loop, as well as DEGs distinguishing the two epithelial loops. Persistence diagrams computed from each method’s latent representations are shown for both cell types. (**C**) Expression patterns of the top-ranked genes contributing to Neuron 46 activation in the ENS context (top) and epithelial context (bottom), as determined by gradient-based attribution.

**Supplemental Figure 7.**
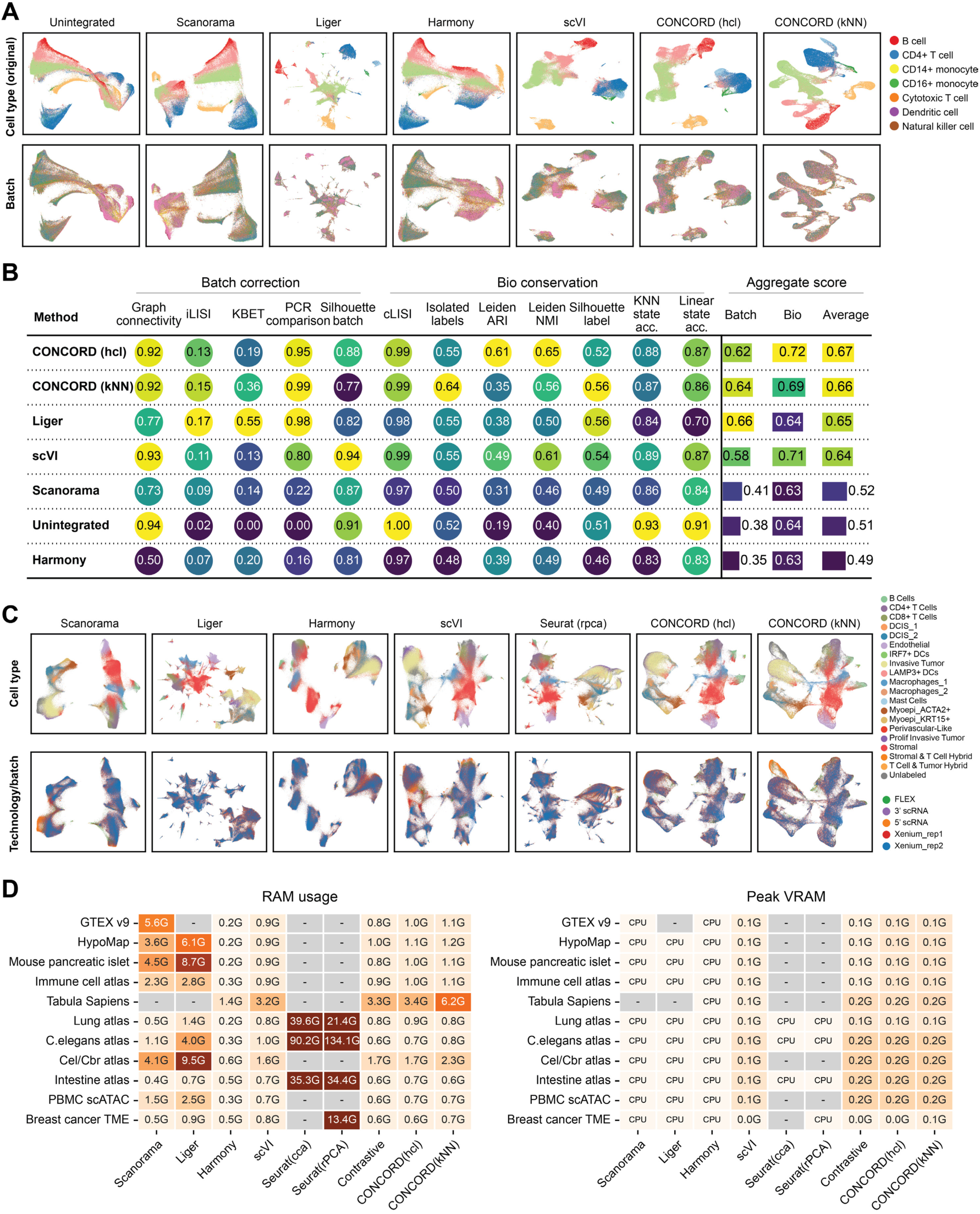
Performance of CONCORD across modalities and scales. (**A**) UMAPs of PBMC scATAC-seq data from unintegrated inputs and from CONCORD and other integration methods, colored by original cell-type annotations and batch. (**B**) Full benchmarking statistics for the PBMC scATAC-seq dataset. (**C**) UMAPs of breast cancer tumor microenvironment data generated by CONCORD and other integration methods, colored by cell type and technology/batch. (**D**) RAM and VRAM usage of different integration methods. For all methods except Seurat, we report ΔRAM (end–start RSS) because Python does not support resetting peak RSS mid-process; for Seurat, we report peak RAM using the *peakRAM* R package. VRAM usage is shown only for GPU-enabled methods. Missing values indicate the method failed to run due to excessive resource requirements or violated assumptions.

**Supplemental Figure 8.**
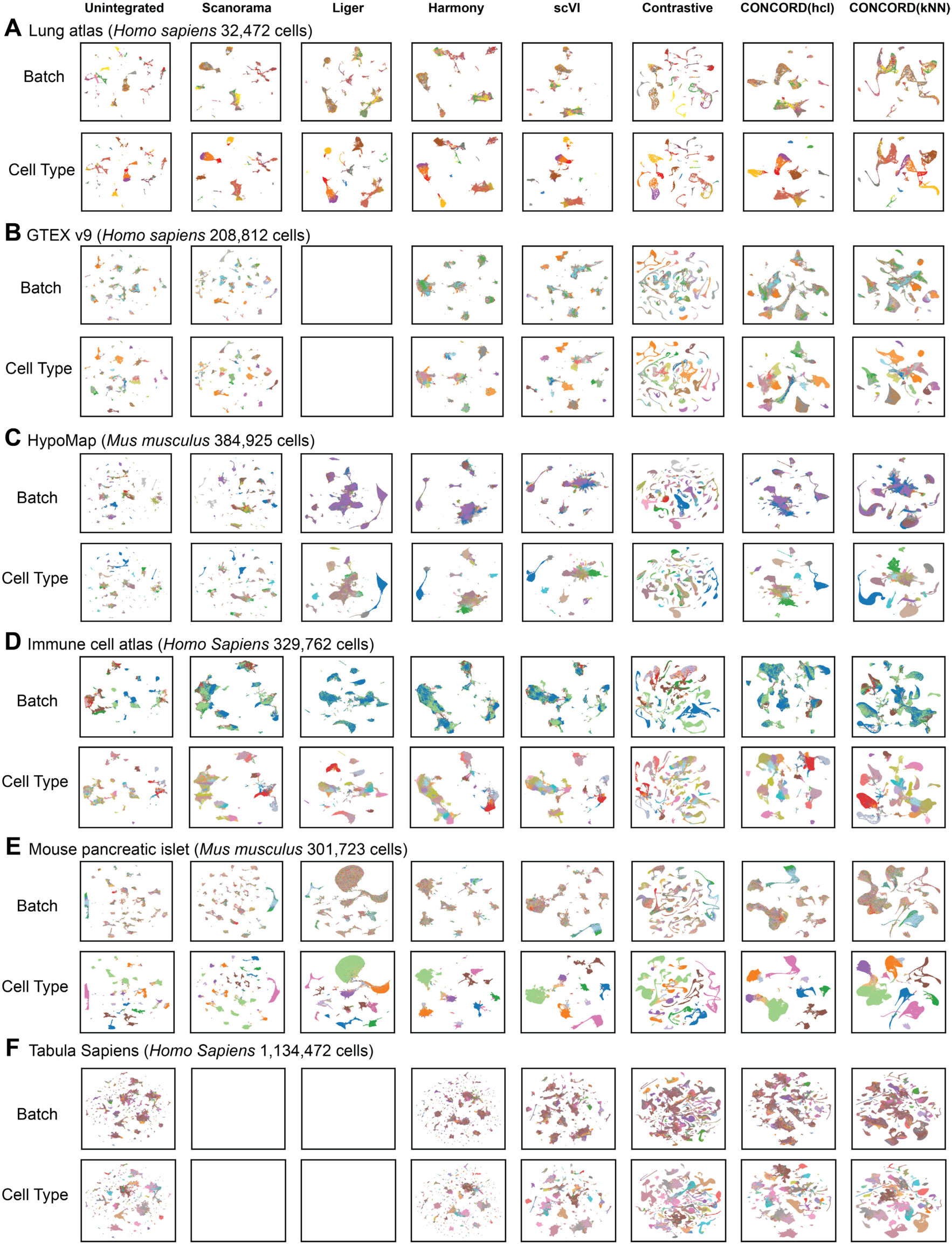
Performance of CONCORD on public human and mouse scRNA-seq datasets. UMAP embeddings derived from each method’s latent space are shown for all datasets and colored by batch and cell type. (A) Lung atlas spanning multiple spatial regions, donors, and two scRNA-seq protocols^33^. (B) GTEX v9: human single-nucleus RNA-seq data from eight tissue types across 16 individuals^81^. (C) HypoMap: single-cell atlas of the murine hypothalamus (∼380k cells) across four assays^82^. (D) Immune cell atlas: human immune cells from 16 tissues and 12 donors^83^. (E) Mouse pancreatic islet: scRNA-seq atlas comprising 56 samples across sex, age, and diabetes models^84^. (F) Tabula Sapiens: human cell atlas of over 1.1M cells from 28 organs of 24 normal human donors^68^.

### Supplementary Tables

**Supplemental Table 1. Benchmarking data-integration methods across diverse simulated structures.** For each structure type (clusters, trajectories, loops, and trees), we simulated varying degrees of batch overlap (full, partial and connected). For the cluster setting, we additionally varied the number of batches and their imbalance. The table reports, for each method and each simulation, scores for topological and geometric preservation, batch correction, and biological-label conservation.

**Supplemental Table 2. Benchmarking data-integration methods across real-world datasets.** For each dataset, we computed batch-correction and biological-label conservation metrics using the scIB^33^ package and probing classifiers. The table reports per-method metric values and summary scores.

